# Engineering anaerobic fungal-bacterial consortia for direct conversion of lignocellulosic biomass into medium-chain fatty acids

**DOI:** 10.1101/2025.08.19.670934

**Authors:** Byung-Chul R. Kim, Elaina M. Blair, Joel P. Howard, Ian M. Gois, Robert Flick, Thomas S Lankiewicz, Stephen Mondo, Jasmyn Pangilinan, Anna Lipzen, Jie Guo, Hope Hundley, Raymond Lee, Jayson Talag, Victoria Bunting, Shanmugam Rajasekar, Kerrie Barry, Igor V. Grigoriev, Michelle A. O’Malley, Christopher E. Lawson

**Affiliations:** Department of Chemical Engineering & Applied Chemistry, University of Toronto, Toronto, ON, Canada; Department of Chemical Engineering, University of California, Santa Barbara, CA, USA; US Department of Energy Joint Genome Institute, Lawrence Berkeley National Laboratory, Berkeley, CA, USA; Arizona Genomics Institute, School of Plant Sciences, University of Arizona, Tucson, AZ, USA; Department of Plant and Microbial Biology, University of California Berkeley, Berkeley, CA, USA; Department of Bioengineering, University of California, Santa Barbara, CA, USA

**Keywords:** Lignocellulosic Biorefinery, Microbiome Engineering, Industrial Biomanufacturing, Division of Metabolic Labor, Anaerobic Fungi, Microbial Chain Elongation, Medium-chain Fatty Acids

## Abstract

Lignocellulosic biomass is a renewable feedstock for sustainable fuels and chemicals, yet industrial conversion remains constrained by carbohydrate solubilization. Inspired by herbivore rumen microbiomes, we engineered an anaerobic fungal-bacterial consortium converting native lignocellulose into medium-chain fatty acids (MCFAs) without pretreatment. Systematic screening identified a newly isolated anaerobic fungus, *Neocallimastix* sp. FC1, in co-culture with *Megasphaera hexanoica* as a top-performing pair, achieving a lignocellulose-to-MCFA yield of 21.0 % (carbon-to-carbon basis) through tight lactate cross-feeding without competition for soluble sugars. Because fungal lactate production rate constrained the growth of *M. hexanoica*, the bacterium reallocated protein from growth toward chain elongation, resulting in increased MCFAs production over butyrate. These results demonstrate that high lignocellulose-to-MCFA conversion by the consortium requires high lactate-producing capability and operating regimes sustaining low lactate concentrations at high flux. Technoeconomic analysis further identifies the cost and yield thresholds required for economically viable deployment, establishing quantitative design targets for pretreatment-free fungal-bacterial lignocellulose upgrading.

## Introduction

The bioconversion of lignocellulosic biomass represents a central opportunity for sustainable fuels and chemicals, yet its industrial deployment remains constrained by how biomass is deconstructed [1–3]. Conventional lignocellulose bioprocesses rely heavily on energy- and capital-intensive upstream operations, including thermochemical pretreatment, external enzyme addition, and intensive mechanical mixing, to render biomass accessible for microbial conversion [1,4]. In practice, process performance, scalability, and economics are governed primarily by the requirements and constraints of upstream biomass deconstruction [3,5,6]. This structural dependence on upstream deconstruction constitutes a fundamental limitation of current lignocellulose biomanufacturing strategies.

We propose a pretreatment-free lignocellulose biomanufacturing platform by engineering anaerobic fungal-bacterial synthetic consortium that leverages direct biological deconstruction (Figure 1). The platform is organized around lactate-based cross-feeding, separating anaerobic fungal biomass deconstruction from downstream bacterial product upgrading, while stabilizing community structure through a defined division of metabolic labour [7,8]. Additionally, routing carbon through lactate, the fungal fermentation end-product and the bacterial substrate, avoids competition for soluble sugars and enables predictable carbon flow across consortium members. This platform reframes lignocellulose processing from a pretreatment-dominated workflow into a biologically integrated, modular biomanufacturing system.

**Figure 1.**
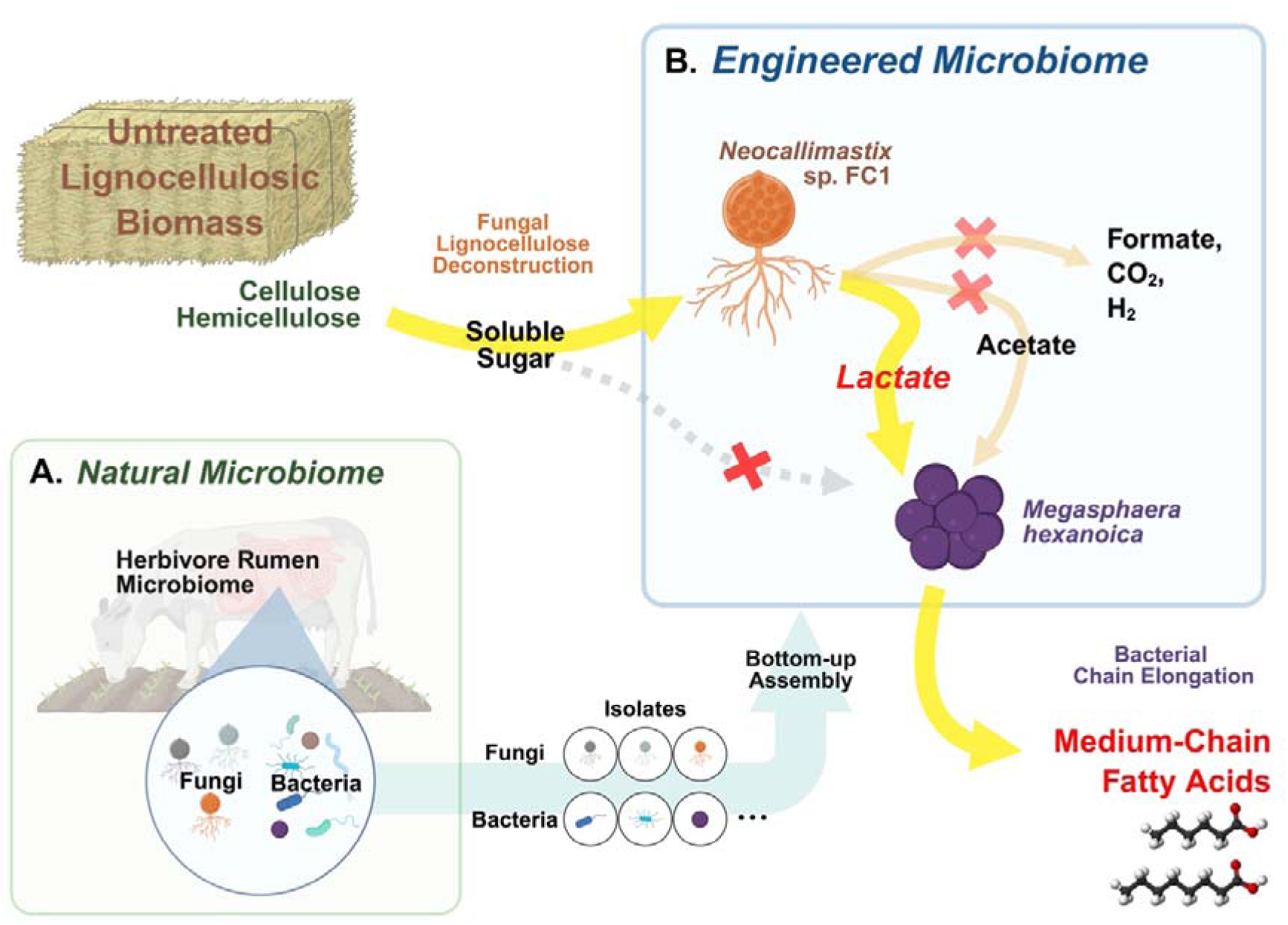
Synthetic microbiome for direct lignocellulose-to-MCFA conversion. Schematic diagram illustrating the engineering of fungal-bacterial synthetic consortia for direct conversion of lignocellulosic biomass into medium-chain fatty acids (MCFAs). **(A)** The bottom-left panel depicts the natural rumen microbiome, where lignocellulose degradation is carried out by a consortium of anaerobic microbes. Microorganisms known to perform target functions are isolated or sourced to establish a curated strain library, which provides the foundation for bottom-up assembly of synthetic fungal-bacterial consortia. **(B)** The top-right panel illustrates an engineered fungal-bacterial consortium that directly converts untreated lignocellulosic biomass into MCFAs. Anaerobic fungi deconstruct lignocellulose and produce lactate and acetate, which are subsequently upgraded by chain-elongating bacteria into MCFAs. The engineered microbiome is organized around lactate-based cross-feeding, avoiding competition for soluble sugars and enabling predictable carbon flow across consortium members. By eliminating alternative carbon sinks and tightening lactate cross-feeding, the platform concentrates flux toward MCFA production. Created with BioRender.com.

The conceptual foundation of this platform draws inspiration from herbivore rumen microbiomes, where lignocellulose deconstruction and fermentation are efficiently integrated. Within these ecosystems, anaerobic fungi play a dominant functional role in lignocellulose deconstruction, constituting only 7-9 % of the rumen microbiota while contributing up to 50 % of fermentable sugar release from ingested plant material [9–11]. This efficiency arises from their rhizoidal growth into intact plant tissues [12] and the coordinated deployment of extensive carbohydrate-active enzyme repertoires within cellulosome-like complexes [13,14]. Consequently, anaerobic fungi facilitate direct biological deconstruction of native plant cell walls, offering a biological alternative to conventional physicochemical biomass processing units.

Following deconstruction, anaerobic fungi perform mixed-acid fermentation, producing primarily formate, acetate, lactate, along with ethanol, hydrogen, and carbon dioxide [15]. As low-molecular-weight, water-soluble compounds produced as a mixed product spectrum, these metabolites are poorly suited as final outputs, as their selective recovery and concentration require energy-intensive separations [16,17]. Therefore, fungal lignocellulose fermentation has typically been coupled with downstream upgrading processes, yielding products such as methane [18–20], butyrate/butanol [21], ethyl acetate/2-phenyl ethanol/isoamyl alcohol [22], and medium-chain fatty acids (MCFAs) [23]. However, the absence of a designated cross-feeding intermediate has led downstream microbes to compete for soluble sugars with anaerobic fungi, preventing clear separation between deconstruction and downstream upgrading and limiting rational control of carbon flow.

To address this gap, we design carbon flow around lactate as a central intermediate, leveraging bacterial chain elongation to convert fungal fermentation products into MCFAs. This upgrading is mediated by reverse β-oxidation (RBO), that couples lactate oxidation to the stepwise elongation of short-chain fatty acids into MCFAs [24]. MCFAs are well suited to scalable fermentation systems, as they combine high product value with efficient recovery and stable process operation. They are high-value commodity chemicals, feedstocks for biofuels in heavy-duty and aviation applications [3,24], and serve as versatile precursors to a wide range of downstream products, including alcohols, esters, methyl ketones, and diols [25,26]. Importantly, their relatively high hydrophobicity enables direct recovery from fermentation broths via membrane-based extraction [27] or pertraction [28], reducing downstream energy demand and mitigating product inhibition. Together, these features position MCFAs as well-suited outputs for scalable biomanufacturing platforms. However, existing MCFA production systems are typically based on either pure culture with narrow substrate scope or open cultures that suffer from competition, instability, and compositional drift, limiting process robustness and controllability [8,29].

Together, this study establishes a pretreatment-free lignocellulose biomanufacturing platform by engineering a defined synthetic fungal-bacterial consortium with lactate-based cross-feeding. The platform is enabled by isolating a high-performing anaero bic fungus and systematically screening fungal-bacterial combinations to identify a co-culture that supports efficient lignocellulose-to-MCFA conversion. By routing carbon exclusively through lactate, the selected consortium avoids direct competition for solu ble sugars, consistent with proteomic evidence indicating lactate-centered metabolism in the bacterial partner. In this configuration, lactate produced by the fungus is rapidly consumed by the bacterial partner, preventing lactate accumulation and promoting elongation to the more desirable C6 and C8 fatty acids. Proteomic analysis further supports this interpretation by showing upregulation of RBO proteins in the bacterial partner under lactate-limited conditions. Techno-economic analysis delineates the process performance targets and cost drivers that will govern advancement of the platform toward economic viability. By integrating strain discovery, mechanistic analysis, and technoec onomic evaluation, this work demonstrates how bottom-up microbiome engineering ca n deliver robust biomanufacturing platforms while identifying actionable levers for sys tematic optimization.

## Results

### Co-culture screening identifies optimal fungal-bacterial partners for MCFAs production

To identify optimal co-cultures for lignocellulose conversion into MCFAs, we first isolated a high-lactate-producing fungal strain, *Neocallimastix* sp. FC1, from a lignocellulose-degrading enrichment culture derived from a cow fecal sample [30]. We sequenced and annotated its genome to support systems-level analysis, which is summarized in Supplementary Table 1. Subsequently, we selected promising strains of anaerobic fungi and chain elongating bacteria based on previously reported biomass degradation and lactate utilization capabilities, respectively, and the availability of genome sequence data to enable systems-level investigation of microbial interactions. This resulted in 3 anaerobic fungal isolates (*Neocallimastix* sp. FC1*, Neocallimastix camerooni* var. *constans,* and *Caecomyces churrovis*) and 7 lactate-utilizing chain elongating isolates (*Caproicibacter fermentans, Caproiciproducens* sp. 7D4C2*, Clostridium luticellarii, Pseudramibacter alactolyticus, Megasphaera cerevisiae, Megasphaera elsdenii,* and *Megasphaera hexanoic*a) that were further screened in monoculture and co-culture to evaluate bioconversion performance (Supplementary Figure 1, 2).

The selected fungal and bacterial isolates were first tested in monoculture to assess their ability to carry out lignocellulose degradation or chain elongation in a chemically defined M2 medium with 10 g/L (1%) milled reed canary grass (1 mm). Fungal monocultures were incubated for seven days, as determined by a preliminary kinetic experiment (Supplementary Figure 3), and 100 mg/L chloramphenicol was supplemented. All three isolates actively degraded reed canary grass, and produced formate, acetate, and lactate as major fermentation products (Figure 2A). In our lignocellulose-to-MCFAs conversion process, lactate acts as a central intermediate in fungal-bacterial cross-feeding, making its titer the key parameter for evaluating the function of fungal isolates. *N*. sp. FC1, a newly isolated fungal strain from this study, exhibited the highest lactate titer, reaching 1.46±0.22 g/L, which was about twice as high as that of *N. camerooni* var. *constans* and *Caecomyces churrovis* (0.76±0.04 and 0.61±0.18 g/L, respectively). The lignocellulose-to-product conversion yield of *N*. sp. FC1 was 49.3 % (carbon-to-carbon basis, see Materials and Methods) after seven days of fermentation and reached 58.1 % after 21 days.

**Figure 2.**
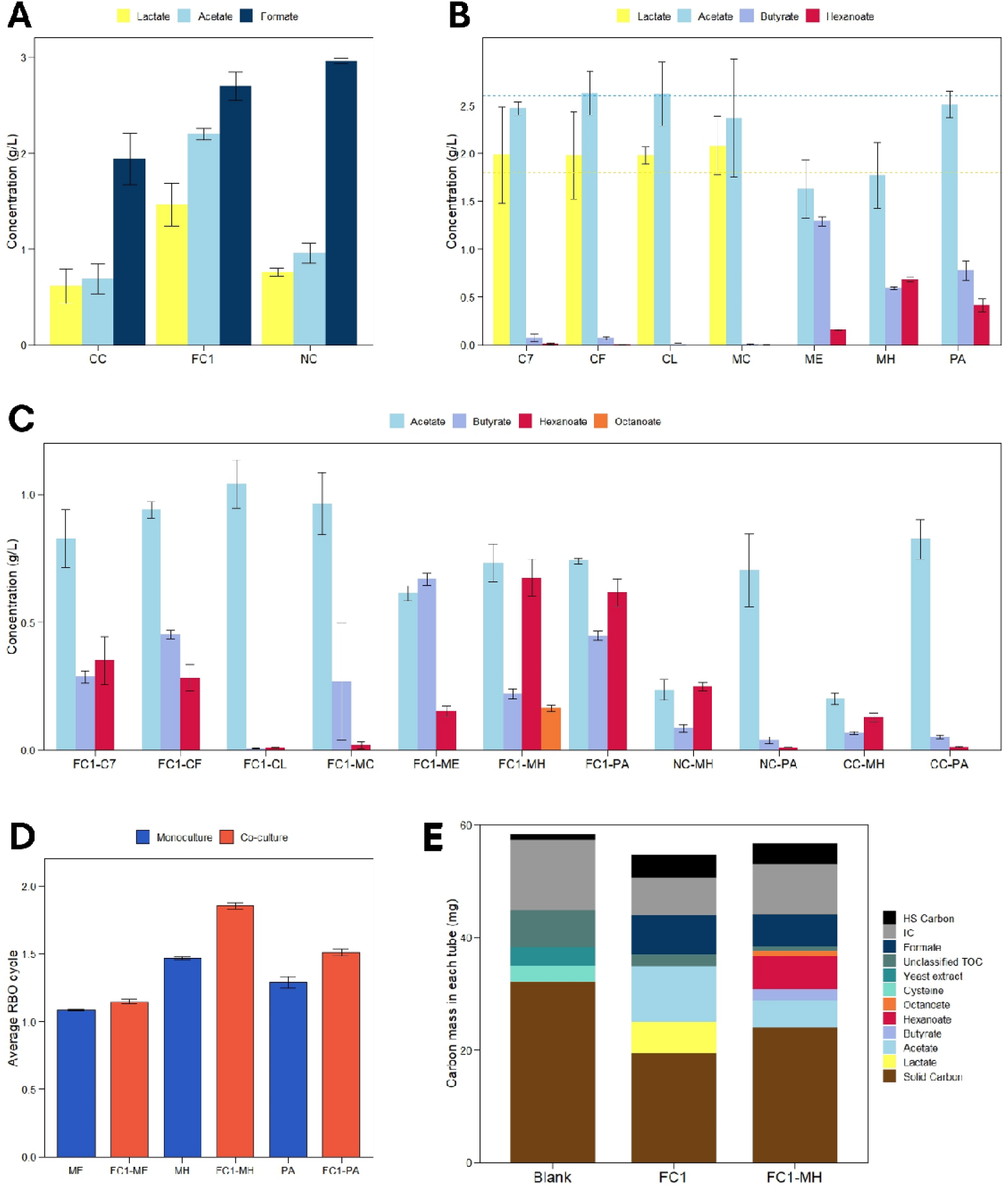
Screening for optimal fungal-bacterial co-cultures. Screening determines an optimal fungal-bacterial co-culture and reveals higher medium-chain fatty acid (MCFA) selectivity in co-cultures than in monocultures. **(A)** Screening of fungal monocultures for lignocellulosic biomass degradation into primary fermentation products. Three fungal isolates were tested: *Caecomyces churrovis* (CC), *Neocallimastix* sp. FC1 (FC1), and *Neocallimastix camerooni* var. constans (NC). **(B)** Screening of bacterial monocultures of seven lactate-utilizing, chain-elongating strains for lactate/acetate-to-MCFA conversion. Strains tested were *Caproiciproducens* sp. 7D4C2 (C7), *Caproicibacter fermentens* (CF), *Clostridium luticellarii* (CL), *Megasphaera cerevisiae* (MC), *Megasphaera elsdenii* (ME), *Megasphaera hexanoica* (MH), and *Pseudoramibacter alactolyticus* (PA). Yellow and sky-blue dotted lines indicate initial concentrations of lactate and acetate, respectively. **(C)** Screening of fungal-bacterial co-cultures to evaluate direct lignocellulose-to-MCFA conversion. Combinations are denoted as “Fungus-Bacterium” (e.g., FC1-MH). **(D)** Average number of reverse β-oxidation (RBO) cycles required for product formation in ME, MH, and PA monocultures and co-cultures with FC1. Butyrate, hexanoate, and octanoate production correspond to one, two, and three RBO cycles, respectively. **(E)** Carbon mass balance for Blank (uninoculated control), FC1, and FC1-MH. Screening data **(A-D)** are shown as means of biological triplicates (error bars indicate standard deviations). Carbon mass balance data **(E)** are shown as means of biological pentaplicates.

The growth of bacterial monocultures was subsequently assessed in the same medium (i.e., M2 medium with reed canary grass) without chloramphenicol but supplemented with yeast extract, lactate, and acetate (1, 1.8, and 2.6 g/L, respectively). Yeast extract was added to support bacterial growth, and the concentrations of lactate and acetate were set to match those produced by *N*. sp. FC1 in monoculture. *M. elsdenii*, *M. hexanoica*, and *P. alactolyticus* produced butyrate and hexanoate, while the four other species didn’t show any clear evidence of growth (Figure 2B). Lactate was completely consumed by all three active strains, whereas acetate utilization differed across strains. *M. elsdenii* and *M. hexanoica* exhibited net consumption of acetate, but *P. alactolyticus* did not. Furthermore, three sister species, *M. cerevisiae*, *M. elsdenii*, and *M. hexanoica* showed diverging results. While *M. cerevisiae* didn’t grow under the condition tested, *M. elsdenii* and *M. hexanoica* showed the highest titer of butyrate (1.29±0.05 g/L) and hexanoate (0.68±0.02 g/L), respectively, among seven chain elongators.

Optimal fungal-bacterial partners for MCFA production were determined by screening co-culture combinations cultivated in M2 medium supplemented with reed canary grass and yeast extract (10 and 1 g/L, respectively) (Figure 2C). The co-culture composed of the top-performing isolates from the fungal and bacterial monoculture screenings, *N*. sp. FC1 and *M. hexanoica*, exhibited the highest lignocellulose-to-MCFAs conversion efficiency (21.0 % carbon-to-carbon yield, see Materials and Methods), and was the only culture that produced octanoate across all screening conditions. The other two bacterial isolates that thrived in the monoculture experiments, *M. elsdenii* and *P. alactolyticus*, also demonstrated robust growth in co-culture. While co-culture containing *M. elsdenii* primarily produced butyrate, the co-culture containing *P. alactolyticus* produced higher levels of hexaonate. Surprisingly, *Caproiciproducens* sp. 7D4C2, *Caproicibacter fermentens*, and *M*. *cerevisiae*, which didn’t grow in monoculture, grew in co-culture. This may reflect fungal-bacterial interactions beyond lactate cross-feeding, although the exact mechanism requires further investigation.

Interestingly, chain elongators (i.e., *M. elsdenii*, *M. hexanoica*, and *P. alactolyticus*) showed higher MCFA specificity when co-cultivated with *N*. sp. FC1 compared to monoculture (Figure 2B, 2C). The average number of RBO cycles involved in product formation was consistently higher in co-cultures with *N*. sp. FC1 than in monocultures (Figure 2D). The increase was more substantial in strains that exhibited higher MCFA specificity in monoculture. To assess whether the enhanced MCFA specificity in co-culture could be attributed to differences in fungal-mediated degradation, we conducted a carbon mass balance analysis comparing monoculture of *N*. sp. FC1 and its co-culture systems with *M. hexanoica* (Figure 2E). This revealed that solid-carbon reduction was lower in the co-culture (25.5 %) compared to the monoculture (39.5 %), likely due to the product (i.e., MCFAs) inhibition on *N*. sp. FC1. Nevertheless, higher MCFA production from a smaller amount of degraded carbon indicated that co-cultivation has higher carbon conversion efficiency toward MCFAs.

### Proteomic analysis highlights lactate-based fungal-bacterial cross-feeding and metabolic shift toward medium-chain fatty acids production in co-culture

To better understand the mechanistic basis of metabolite cross-feeding and its link to improved MCFA production in co-culture, we performed comparative proteomic analysis of monoculture and co-culture systems (Figure 3A, Supplementary dataset 1). This allowed us to establish functional baselines for each organism. In *N*. sp. FC1, malic enzyme, pyruvate formate lyase, and xylose isomerase (Enzymes 17, 18, and 7 in Figure 3A, respectively) were the most abundant proteins (Supplementary Table 2). Malic enzyme and pyruvate formate lyase are key enzymes of the hydrogenosome, a mitochondria-like organelle of anaerobic fungi specialized in producing ATP and H_2_ [15]. This finding is consistent with the observed active production of acetate and formate (Figure 2A). The abundance of xylose isomerases also highlights the potential of anaerobic fungi as a key biocatalyst for lignocellulose-based refineries, given that the lack of efficient industrial xylose-fermenting microbes limits hemicellulose utilization [31,32]. In the *M. hexanoica*, electron transfer flavoprotein, enoyl-CoA hydratase, and pyruvate:ferredoxin oxidoreductase (Enzymes 32, 31, and 28 in Figure 3A, respectively), all key enzymes for chain elongation, were the most abundant proteins (Supplementary Table 2), underscoring the metabolic specialization of this strain.

**Figure 3.**
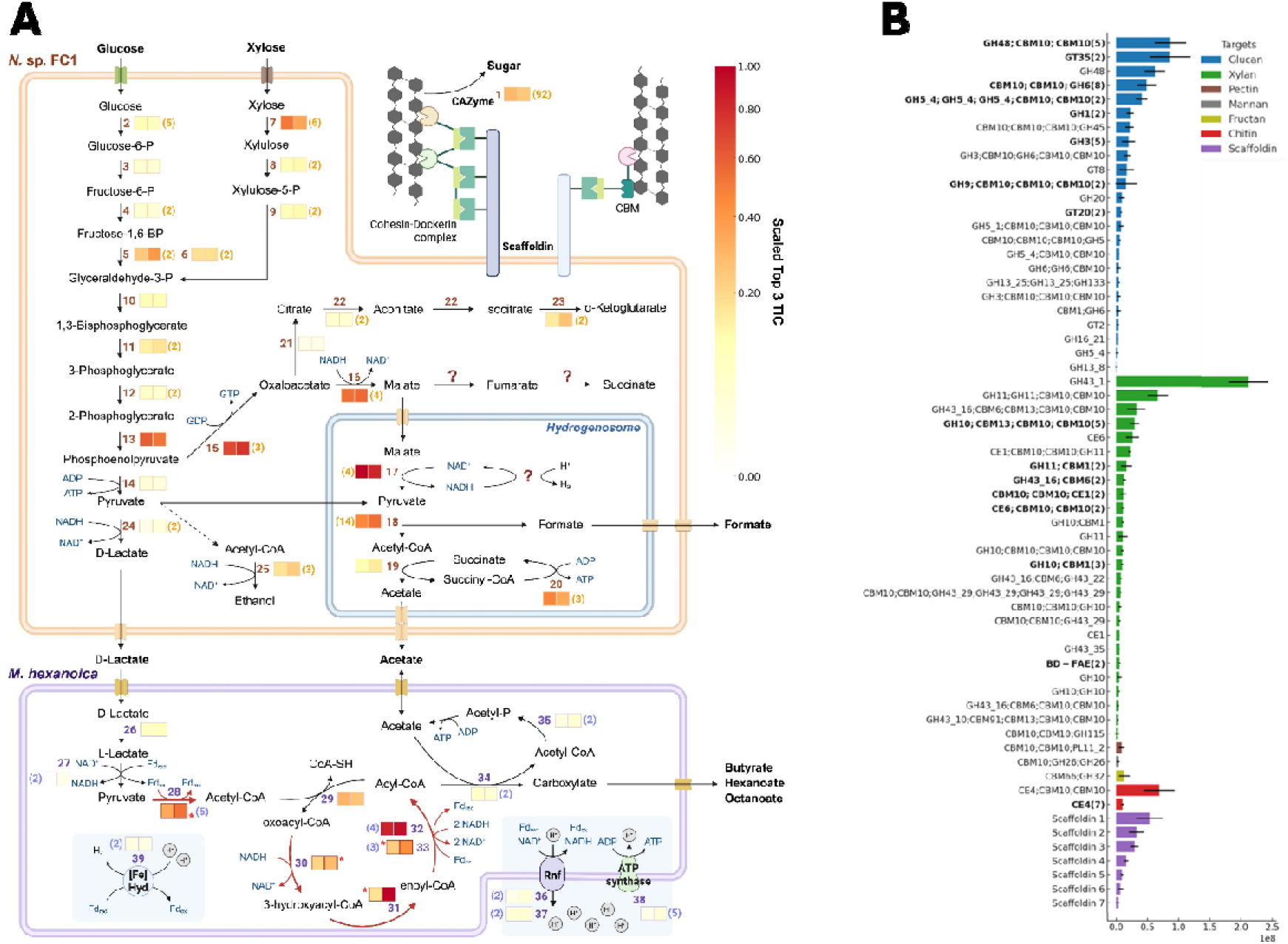
Fungal and bacterial proteomes. Differential proteomics comparing monoculture and co-culture of *Neocallimastix* sp. FC1 and *Megasphaera hexanoica*. **(A)** Integrated metabolic pathway map and differential proteomics of *N*. sp. FC1 (top orange box) and *M. hexanoica* (bottom purple box). Each reaction is accompanied by paired TIC (Top 3 Total Ion Current value) boxes indicating the average of scaled enzyme abundance in monoculture (left) and co-culture (right). For enzymatic steps with multiple detected proteins, only the one with the highest top 3 TIC value was visualized, and the total number of proteins corresponding to that step is indicated in parentheses (light orange for *N*. sp. FC1, light purple for *M. hexanoica*). The color of each TIC box reflects enzyme abundance, normalized to the highest top 3 TIC value observed within each organism: monoculture malic enzyme (enzyme 17) for *N*. sp. FC1 and co-culture enoyl-CoA hydratase (enzyme 31) for *M. hexanoica*. Reactions that were significantly upregulated in co-culture (FDR < 0.05) are marked with red arrows, and a red star symbol is placed next to the corresponding TIC boxes. Created with BioRender.com. **(B)** Top 3 TIC values of carbohydrate-active enzyme (CAZyme) and scaffoldin proteins in *N*. sp. FC1 monoculture. CAZymes with identical domains in the same order are shown in bold, with the number in parentheses indicating the number of proteins sharing that domain architecture. Proteins were classified according to their predicted target substrates (i.e., glucan, xylan, pectin, mannan, fructan, chitin, scaffoldin). Data represent the average of biological quadruplicate, with error bars indicating standard deviations.

Although this study analyzed only intracellular proteins, a diverse set of CAZymes was identified in *N*. sp. FC1 (Figure 3B). A broad array of endoglucanases, exoglucanases, and β-glucosidases (e.g., GH1, GH3, GH5, GH6, GH9, GH13, GH45, and GH48), as well as hemicellulases (e.g., GH10, GH11, GH43, and CE1), were identified (Supplementary dataset 1). Most of the CAZymes contained carbohydrate-binding domains (CBMs), such as CBM10, CBM6, and CBM13, which are known to enhance substrate affinity and synergistic catalysis [33]. In anaerobic fungi, the frequent occurrence of CBM10 is particularly noteworthy, as it may also function as a dockerin-like domain that anchors catalytic subunits to scaffoldins [34]. Indeed, a scaffoldin protein was detected, strongly supporting the presence of a cellulosome-like enzymatic complex in *N*. sp. FC1. This, together with the presence of proteins harboring diverse domain combinations, points to a system that combines broad catalytic capability with specificity toward particular substrates (Figire 3B, Supplementary dataset 1). For example, a single protein sequence contained both GH3 and GH6 domains along with three CBM10 domains, while another protein sequence included GH43_16 together with three distinct CBM domains (CBM6, CBM13, and CBM10). GH43_29 was also detected in four tandem repeats, together with two CBM10 domains. Together, these features may facilitate efficient lignocellulose breakdown and suggest an important role for *N*. sp. FC1 in metabolic exchange within the co-culture.

Metabolite cross-feeding between *N*. sp. FC1 and *M. hexanoica* under co-culture cultivation was predominantly based on D-lactate. *N*. sp. FC1 solely produced D-lactate but not L-lactate, in agreement with genomic evidence showing the presence of a D-lactate dehydrogenase gene and not a gene for L-lactate dehydrogenase. However, the abundance of lactate racemase and L-lactate dehydrogenase in *M. hexanoica* did not differ significantly between monoculture and co-culture (Figure 3A, Supplementary Table 2). The absence of sugar transporters (e.g., PTS subunits IIABC) and metabolic enzymes (e.g., hexokinase, xylose isomerase) in *M. hexanoica* proteome, together with undetectable sugar levels in fungal monoculture, suggests minimal sugar-mediated cross-feeding. Interestingly, enzymes involved in riboflavin biosynthesis (6,7-dimethyl-8-ribityllumazine synthase, Supplementary Table 2) were downregulated in *M. hexanoica* under co-culture, suggesting potential cross-feeding of trace micronutrients. Such micronutrient transfer may have facilitated the growth of strains that failed to grow in monoculture (*Caproiciproducens* sp. 7D4C2, *Caproicibacter fermentens*, and *M. cerevisiae*; Figure 2B, 2C) under co-culture, pointing to fungal-bacterial interactions beyond lactate cross-feeding. In addition, *M. hexanoica* proteins related to nitrogen metabolism exhibited marked shifts in abundance under co-culture, suggesting restricted access to nitrogen source due to competition with the fungal partner (Supplementary Note 1, Supplementary Table 2).

Differential proteomic analysis between monoculture and co-culture conditions revealed that the enhanced production of hexanoate and octanoate over butyrate in co-culture was driven by changes in the bacterial proteome. While *N*. sp. FC1 proteome showed no statistically significant differences, *M. hexanoica* exhibited a marked upregulation of chain elongation pathways under co-culture. All enzymes involved in the RBO cycle were upregulated, except acetyl-CoA C-acetyltransferase (Enzyme 29 in Figure 3A) which catalyzes highly thermodynamically constrained reaction [35] (ΔG°1 = +25.0 kJ/mol for C4 metabolism). The unchanged expression level of acetyl-CoA C-acetyltransferase suggests that this step functions as a strict thermodynamic bottleneck, where additional enzyme investment cannot meaningfully increase flux [35]. In contrast, enoyl-CoA hydratase (Enzyme 31 in Figure 3A), the only other enzyme in the RBO cycle that catalyzes a thermodynamically unfavorable reaction (Enzyme 31 in Figure 3A, ΔG°1 = +3.3 kJ/mol, for C4 metabolism), was 5.9-fold more abundant in co-culture compared to monoculture (Figure 3A), suggesting increased thermodynamic pressure on chain elongation. Meanwhile, the terminal enzyme CoA-transferase (Enzyme 34 in Figure 3A) did not exhibit a significant difference between monoculture and co-culture. The increased ratio of RBO cycle enzymes to the terminal enzyme under co-culture condition reflects an expression profile that promotes greater RBO cycle iterations per product, consistent with the elevated production of C6 and C8 (Figure 2D).

### Kinetic analysis elucidates the role of cross-feeding intermediates in driving chain elongation

Given the central role of lactate as a cross-feeding intermediate, we conducted a kinetic experiment on the co-culture of *N*. sp. FC1 and *M. hexanoica* to assess whether lactate is transiently accumulated or consistently depleted. Lactate was not detected at any point during the 10-day incubation, indicating that it is not only the main driver of chain elongation but also the principal limiting metabolite in lignocellulose-to-MCFA conversion. We propose that this persistent lactate scarcity triggered increased chain elongation in the co-culture condition [36], supported by the observed increase in RBO enzyme expression. This proteome allocation shift is consistent with a compensatory strategy in which cells upregulate enzyme abundance to maintain metabolic flux under substrate-limited conditions [37].

We also found that another cross-feeding metabolite, acetate, governed the MCFAs production kinetics in co-culture conditions. The dynamics of MCFAs production by *M. hexanoica*, which actively consumed acetate during chain elongation, were decoupled from those of fungal growth and metabolite release. The accumulated headspace pressure, which correlates with fungal proliferation and lactate/acetate production, increased rapidly in the first five days (Supplementary Figure 3) [38]. However, the MCFAs production pattern of *M. hexanoica* was progressively distributed with longer-chain MCFAs increasingly produced during the latter half of the incubation (Figure 4A, 4C). In contrast, chain elongation kinetics of *P. alactolyticus*, which did not assimilate acetate, closely resembled the kinetics of fungal proliferation and assumed lactate production (Figure 4B, 4C). These dynamics imply that *P. alactolyticus* operates under pseudo-first-order kinetics on lactate during lactate-scarcity, with chain elongation flux responding directly to transient substrate supply from fungal metabolism. Together, these findings suggest that acetate utilization capacity may play an important role in shaping the temporal organization of chain elongation dynamics in cross-feeding systems.

**Figure 4.**
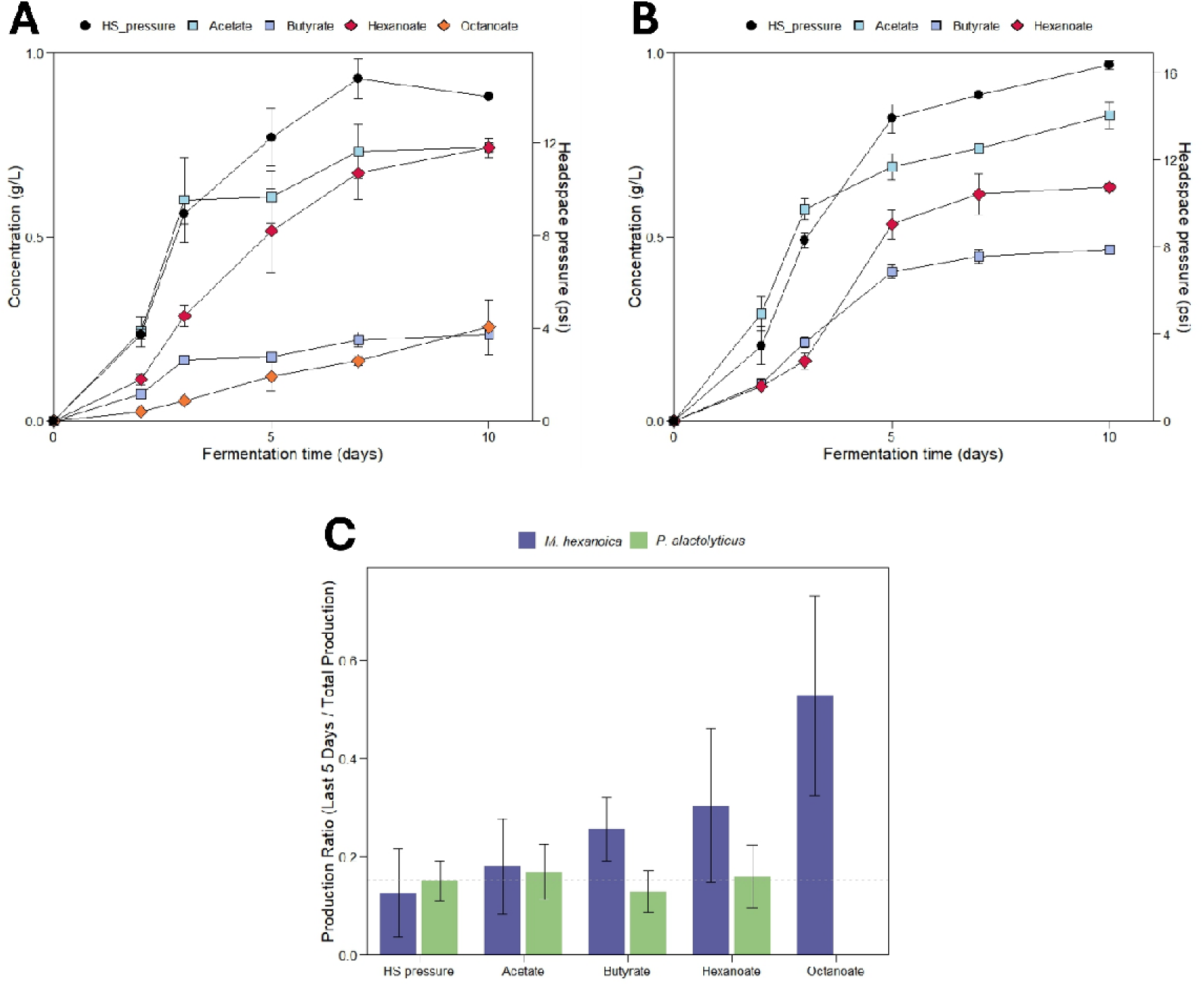
Chain elongation kinetics. Kinetic analysis of fungal-bacterial co-cultures demonstrates tight lactate cross-feeding and differences in chain elongation dynamics depending on acetate involvement. **(A-B)**, Kinetics of lignocellulose-to-product conversion in co-cultures of *Neocallimastix* sp. FC1 with *M. hexanoica* (**A**) and *P. alactolyticus* (**B**). **(C)** Contribution of each product during the day 5-10 relative to total production in co-cultures of *Neocallimastix* sp. FC1 with *M. hexanoica* or *P. alactolyticus*. All data represent the average of biological triplicates, with error bars indicating standard deviations.

### Techno-economic analysis suggests cost and yield benchmarks for scalable fungal-bacterial CBP systems

Technoeconomic analysis was performed to validate the potential of anaerobic fungal-bacterial consortia as a profitable biorefinery platform and outline design criteria to achieve its practical implementation. This analysis was conducted using the open-source TEA platform BioSTEAM [39] and parameterized based on prior experiments and simple assumptions (Supplementary Note 2, Supplementary Table 3). The model comprises media preparation, fermentation, separation, and distillation (Figure 5A). An initial analysis, based on experimental results and unoptimized conditions from this study, revealed economic infeasibility. This primarily arose from the high medium cost per product ($127.1 USD/kg_MCFAs), with cysteine (49.7 %), yeast extract (39.4 %), and buffering agents (carbonate and phosphate buffer, 9.5 %) accounting for the majority of expenses (Figure 5B). The low MCFA titer further increased the cost per product, as the fixed medium cost was allocated across a smaller yield, which is constrained by the low lignocellulose loading (10 g/L) and MCFA yield (0.1 g_MCFAs/g_grass). Sensitivity analysis evaluated the influence of key process design criteria on IRR (Figure 5C) and utility cost (Figure 5D), with medium cost, MCFAs yield, and RCG input emerging as the most influential factors. Therefore, we analyzed profitability across different medium costs and MCFA yields (Figure 5E and 5F for hexanoate and octanoate, respectively) under higher lignocellulose loading (100 g/L) to propose economically feasible engineering design criteria. The results suggest economically feasible targets for medium cost and MCFA yield, represented by the internal rate of return (IRR) contour lines (from left to right, representing IRR levels of 0.1, 0.15, and 0.2, respectively; see Figure 5E and 5F). Overall, minimizing medium costs and intensifying the process play a crucial role in transitioning this early-stage bioprocess into a scalable and economically viable biomanufacturing system.

**Figure 5.**
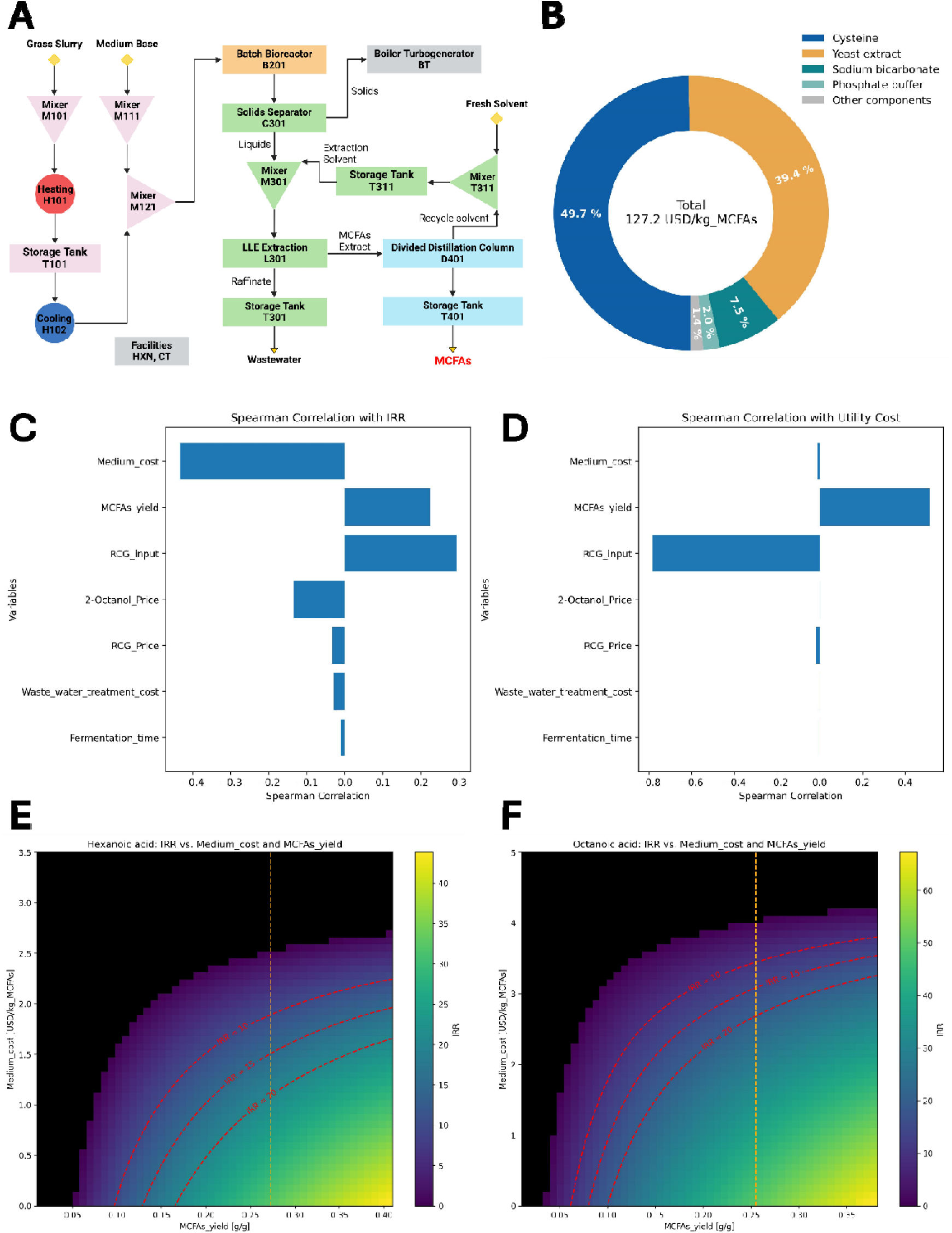
Techno-economic analysis. Techno-economic analysis of a fungal-bacterial CBP system for lignocellulose-to-MCFA conversion. **(A)** Process flow diagram of the fungal-bacterial CBP system for lignocellulose-to-MCFA conversion, including media preparation, fermentation, separation, and distillation. **(B)** Pie chart showing the relative cost contribution of medium components for bioreactor cultivation. **(C-D)** Sensitivity analysis identifying key process parameters influencing economic performance, measured by internal rate of return (IRR, **C**) and utility cost (**D**). **(E-F)** Simulated IRR as a function of medium cost (y-axis) and MCFA yield (x-axis) for hexanoic acid (**E**) and octanoic acid (**F**) production. Red dashed contours represent IRR thresholds (0.1, 0.15, 0.2). A yellow vertical dashed line marks the theoretical maximum MCFA yield for a two-member co-culture of an anaerobic fungus and a chain elongating bacterium. The x-axis upper bound corresponds to the theoretical maximum MCFA yield under a CO_2_ recovery scenario in which homoacetogens and ethanol-utilizing chain elongating bacteria are added to the synthetic consortium.

## Discussion

This study demonstrates a microbiome engineering framework to construct a synthetic fungal-bacterial consortium for pretreatment-free conversion of lignocellulosic biomass into MCFAs. Through bottom-up assembly, we show that efficient lignocellulose-to-MCFA conversion can be achieved by pairing top-performing fungal (*N*. sp. FC1, isolated from this study) and bacterial strains (*M. hexanoica*). This underscores strain selection as a primary design variable in engineered microbiomes which can be systematically accelerated through culturomics [8,40]. Within this consortium, lignocellulose-to-MCFA conversion was enabled by a lactate-centered cross-feeding between anaerobic fungi and chain-elongating bacteria preventing direct competition for soluble sugars. This result is consistent with prior report showing that *M. hexanoica* rapidly consumes lactate while largely bypassing glucose [36]. Additionally, lactate and acetate played distinct functional roles: lactate served as the primary fuel for chain elongation by sustaining flux through RBO, whereas acetate modulated the timing of product formation, with higher acetate involvement associated with delayed accumulation of hexanoate and octanoate. Notably, preferential production of hexanoate and octanoate under lactate-scarce conditions indicates that maintaining low lactate levels while maximizing RBO flux can enhance lignocellulose-to-MCFA productivity. From an industrial perspective, this product shift is advantageous, as longer-chain fatty acids exhibit higher market value and greater compatibility with product recovery than butyrate [28]. Moreover, because acetate represents a relatively oxidized cross-feeding intermediate, limiting its contribution to chain elongation further favors the formation of more reduced fatty acids [35]. These findings highlight the importance of securing homo-lactic acid fungi to ensure consistent lactate supply while minimizing acetate input that may otherwise divert carbon away from MCFAs.

The preferential production of hexanoate and octanoate under lactate-limited co-culture conditions demonstrates a generalizable design principle for biomanufacturing systems targeting reduced chemical products. Rapid depletion of lactate, the primary carbon and electron donor for RBO, indicates that electron donor demand by chain-elongating bacteria exceeded fungal supply, imposing a growth-limited regime. Such growth limitation is known to shift metabolic investment away from biosynthesis and toward energy-conserving fermentative pathways in anaerobic microbes [41], a physiological state supported here by increased abundance of RBO-associated enzymes in *M. hexanoica* during co-culture. Under these conditions, reduced anabolic demand lowers electron consumption for biomass synthesis, creating a surplus of reducing equivalents that must be rebalanced through metabolism. This surplus reducing power was channeled into the formation of more reduced fermentation end-products. Accordingly, the system favored accumulation of hexanoate and octanoate, which require additional elongation steps and greater enzymatic investment than butyrate [35] yet retain a larger fraction of reducing equivalents in the final products. Similar shifts toward more reduced products have been reported across diverse anaerobic systems, including the transition from acidogenesis to solventogenesis in *Clostridium acetobutylicum* as growth declines in batch culture [42] and the redirection of carbon flux from acetate to ethanol in *Clostridium autoethanogenum* during electron donor-limited chemostat operation at high biomass density [43]. Together, these observations suggest that deliberately operating anaerobic bioprocesses within a controlled growth-limited regime can promote redox rebalancing through reduced product formation, maximizing carbon and electron retention in target chemicals.

The technoeconomic analysis further supports the economic feasibility of this platform and clarifies the cost-yield conditions required for profitable implementation. Achieving this potential will require advances across three key domains: medium cost reduction, metabolic optimization, and process intensification. The unique ability of anaerobic fungi to deconstruct lignocellulosic biomass without extensive pretreatment or enzymatic hydrolysis enables high solid loadings [44], offering a practical strategy to medium cost reduction and process intensification. Additional cost savings can be achieved by replacing expensive medium components such as cysteine, buffer salts, and yeast extract with renewable alternatives or intensifying microbial cross-feeding [45,46]. Metabolic optimization strategies, such as enhancing lactate production through hydrogenosomal regulation [44,47], refining MCFA selectivity in chain-elongating bacteria [48], and expanding functional guilds (e.g., homoacetogens, ethanol-based chain elongators) [8], could significantly boost productivity and carbon efficiency. For example, incorporating homoacetogens and ethanol-based chain elongators increases the theoretical maximum MCFA yields of hexanoate and octanoate from 27.3 and 25.5 % (yellow vertical dashed line in Figure 5E, 5F) to 41.0 and 38.2 % (the upper bound of x-axis in Figure 5E, 5F), respectively. Lastly, process intensification approaches, including high-solids continuous-flow culturing [49,50], solid-state fermentation [51], and membrane-based extraction technologies [28], offer promising routes to improve volumetric productivity and mitigate product inhibition.

### Concluding remarks

This study establishes a defined anaerobic fungal-bacterial consortium as a pretreatment-free platform for lignocellulose biomanufacturing, enabling direct conversion of native plant biomass into MCFAs. By systematically screening defined fungal-bacterial pairings, an optimal consortium is identified, achieving conversion of 21 % of lignocellulosic carbon into MCFAs. The results further identify operationally controllable factors, including lactate availability and acetate utilization capacity, as key levers shaping product selectivity and kinetics, providing practical guidance for tuning system behavior. In parallel, technoeconomic analysis translates the demonstrated platform performance into quantitative benchmarks by delineating the cost and yield targets required for economically viable deployment, clarifying where future optimization efforts are most impactful.

Taken together, these findings position fungal-bacterial consortia as a biological front end for lignocellulose-based biorefineries, channeling lignocellulosic biomass into MCFAs that provide a common starting point for diverse downstream biological or catalytic upgrading strategies [3,25]. More broadly, the work shifts lignocellulose biomanufacturing from a pretreatment-centered challenge toward a system integration problem that becomes tractable through systematic organization of biological functions and process design.

## Materials and methods

### Strains and routine cultivation

In this study, two anaerobic fungal strains previously isolated in earlier experiments (*Neocallimastix camerooni* var. constans [52], and *Caecomyces churrovis* [19]) and newly isolated strain (*Neocallimastix* sp. FC1) were used. Isolation, cryopreservation, and subsequent revival of the fungal strains were performed following established protocols for anaerobic gut fungi [9]. *Neocallimastix* sp. FC1 was obtained from lignocellulose-degrading enrichment culture derived from a cow fecal sample [30]. Fungal strains were first revived in 160 mL serum bottles containing 50 mL MC medium supplemented with 10□g/L of milled reed canary grass (1□mm particle size). They were then routinely subcultured every 3 days in M2 medium supplemented with chloramphenicol (100□mg/L) and reed canary grass (10□g/L), using Hungate tubes with a 1□mL inoculum and a 10□mL working volume. Reed canary grass was provided by the U.S. Department of Agriculture, Agricultural Research Service, U.S. Dairy Forage Research Center [21].

Lactate-utilizing chain elongating bacterial strains (*Caproicibacter fermentans* DSM 107079, *Caproiciproducens* sp. 7D4C2 DSM 110548, *Clostridium luticellarii* DSM 29923, *Pseudramibacter alactolyticus* DSM 3980, *Megasphaera cerevisiae* DSM 20461, *Megasphaera elsdenii* DSM20460, and *Megasphaera hexanoica* DSM 106893) were purchased from the German Collection of Micro-organisms and Cell Cultures (DSMZ). Bacterial cultures were revived in ATCC 2107 medium supplemented with sodium lactate (10□g/L) but without soluble starch and dextrose. They were subcultured under the same condition after 4 days of cultivation, by transferring 0.02□mL of preculture into Hungate tubes with 10□mL working volume. For both fungal and bacterial cultures, bottles and tubes were prepared and sealed within a glovebox to maintain strict anaerobic conditions prior to autoclaving. Inoculated cultures were subsequently incubated at 39 °C in a water bath.

### Genomic DNA extraction, sequencing, and genome annotation of *Neocallimastix* sp. FC1

Fungal cell mats of *Neocallimastix* sp. FC1 were harvested after four days of growth on sorghum in three different media: MC, M2, and M2 supplemented with 1□g/L yeast extract. The samples were shipped on dry ice to the Arizona Genome Institute for genomic DNA and total RNA extraction. High-quality genomic DNA was extracted at the Arizona Genomics Institute using a modified cetyl trimethylammonium bromide (CTAB) protocol [53]. Briefly, the frozen fungal biomass was cryo-ground using mortar and pestle with liquid nitrogen and subjected to extraction with CTAB buffer for 1 hour at 50 °C. After centrifugation, the supernatant was purified by two repeated extractions with chloroform-isoamyl alcohol (24:1). The upper aqueous phase was recovered and mixed with 3□M sodium acetate (10 % v/v), and genomic DNA was precipitated with isopropanol. The DNA pellet was collected by centrifugation, washed with 70□% ethanol, air-dried for 20□minutes, and dissolved in Tris-EDTA buffer (10 mM Tris, pH 8, 1 mM EDTA) at room temperature followed by RNAse treatment. Subsequently, DNA purity was assessed with a NanoDrop spectrophotometer, concentration was measured with the Qubit HS kit (Invitrogen), and size was validated with the Femto Pulse System (Agilent). The purified genomic DNA of *Neocallimastix* sp. FC1 was shipped to the Joint Genome Institute (JGI) for sequencing and annotation.

For genome sequencing, 1500□ng of genomic DNA was sheared to approximately 10□kb using either a Megaruptor® 3 (Diagenode) or a g-TUBE (Covaris). The sheared DNA was first treated with exonuclease to remove single-stranded ends, followed by DNA damage repair, end-repair, and A-tailing using reagents from the SMRTbell Express Template Prep Kit 2.0 (PacBio). Barcoded overhang adapters from the SMRTbell® Barcoded Adapter Plate 3.0 (PacBio) were then ligated to the prepared DNA fragments, and the resulting libraries were purified using AMPure® PB Beads (PacBio). The purified libraries were pooled based on the required number of reads perd sample to ensure sufficient sequencing coverage. Pooled libraries were size selected using 0.75□% agarose gel cassettes with Marker S1 and the High Pass protocol on a BluePippin system (Sage Science). A sequencing primer was annealed to the SMRTbell templates, and the polymerase was bound using the Sequel II Binding Kit 2.0 (PacBio). Sequencing was performed on a PacBio Sequel IIe platform using SMRT Link v10.2, 8M v1 SMRT cells, and Sequel II Sequencing Kit v2.0, with 1×1800 sequencing movie run times. Circular consensus sequencing (CCS) reads were filtered to remove artifacts and assembled using Flye v2.9-b1768 in HiFi mode [54]. The resulting draft genome was polished using two iterations of Racon v1.4.13 and annotated through the JGI annotation pipeline using assembled transcriptome data [55].

### Total RNA extraction, sequencing, and transcriptome assembly of *Neocallimastix* sp. FC1

Total RNA was extracted at the Arizona Genomics Institute using the Qiagen RNeasy® Mini Kit, following the manufacturer’s protocol described in the RNeasy® Mini Handbook under the section “Purification of total RNA from plant cells and tissues and filamentous fungi”. An on-column DNase digestion step was included to remove residual genomic DNA. Fungal cells were lysed by grinding in liquid nitrogen, and homogenization was performed using a QIAshredder. RNA was eluted in 50□μL of RNase-free water, then re-applied to the column to concentrate the final RNA solution. RNA concentrations exceeded 25□ng/μL as measured by the Invitrogen Qubit 2.0 fluorometer. Integrity was assessed using either the Agilent 2200 TapeStation or 2100 Bioanalyzer, with all samples showing RIN or RINe scores above 7. The purified total RNA from *Neocallimastix* sp. FC1 was then shipped to the Joint Genome Institute (JGI) for sequencing and assembly.

For Illumina RNA-Seq, mRNA was isolated from an input of 1000 ng of total RNA with oligo dT magnetic beads and fragmented to 300 bp - 400 bp with divalent cations at a high temperature. Using TruSeq stranded mRNA kit (Illumina), the fragmented mRNA was reverse transcribed to create the first strand of cDNA with random hexamers and SuperScript™ II Reverse Transcriptase (Thermo Fisher Scientific) followed by second strand synthesis. The double stranded cDNA fragments were treated with A-tailing, ligation with NEXTFLEX UDI Barcodes (PerkinElmer) and enriched using 8 cycles of PCR. The prepared libraries were quantified using KAPA Biosystems’ next-generation sequencing library qPCR kit and run on a Roche LightCycler 480 real-time PCR instrument. Sequencing of the flow cell was performed on the Illumina NovaSeq sequencer using NovaSeq XP V1.5 reagent kits, S4 flowcell, following a 2×151 indexed run recipe. RNA-Seq reads were filtered and trimmed for quality and contamination using BBDuk (https://sourceforge.net/projects/bbmap/) and assembled using Trinity v2.12.0 [56].

For PacBio Iso-Seq, 500 ng of total RNA was used to synthesize full-length cDNA and enriched using 12-20 cycles of PCR with NEBNext® Single Cell/Low Input cDNA Synthesis & Amplification Module (New England BioLabs). The amplified cDNA was purified (or size-selected) with AMPure® PB Beads (PacBio). Libraries were pooled based on the required amount of reads per sample, ensuring each sample received adequate coverage for analysis. The pooled libraries were treated with DNA damage repair enzyme mix, end-repair/A-tailing mix and ligated with barcoded overhang adapters from SMRTbell Barcoded Adapter Plate 3.0 (PacBio) using SMRTbell Express Template Prep Kit 2.0 (PacBio). PacBio Sequencing primer was then annealed to the SMRTbell template library and sequencing polymerase was bound to them using Sequel II Binding kit 2.0. The prepared SMRTbell template libraries were then sequenced on a Pacific Biosystems’ Sequel IIe sequencer using SMRT Link 12.0, 8M v1 SMRT cells, and Version 2.0 sequencing chemistry with 1×1800 sequencing movie run times. Polished Circular Consensus Sequence (CCS reads) were produced from Pacbio subreads using pbccs v. 4 (ccs --min-passes 3 --min-snr 4 --max-length 21000 --min-length 10 --min-rq 0.98) as described in the official PacBio Iso-Seq analysis pipeline (https://github.com/PacificBiosciences/IsoSeq), classified as full length based on 5’ and 3’ primers and polyA tail, trimmed for polyA tails, and clustered all the full length CCS reads to produce a single consensus sequence per cluster, following the Iso-Seq protocol [57].

### Screening and kinetics experiments

To identify optimal fungal-bacterial pairs for MCFA production, we conducted screening experiments of monocultures and co-cultures using a M2 medium supplemented with 10□g/L (1□%) milled reed canary grass (1□mm particle size). Fungal monocultures were grown for 7 days with 100□mg/L chloramphenicol, with the incubation period determined based on a preliminary kinetic experiment (Supplementary Figure 3). Bacterial monocultures were subsequently assessed under the same medium condition without chloramphenicol, but with additional supplementation of yeast extract, lactate, and acetate (1, 1.8, and 2.6□g/L, respectively) to support growth and mimic the metabolite profile produced by *N*. sp. FC1 in monoculture. Co-cultures were cultivated in the same medium with 1□g/L of yeast extract but without lactate and acetate to test direct conversion of reed canary grass into MCFAs. All cultures were grown in Hungate tubes with 10□mL working volume, and all experiments were conducted in biological triplicate. For inoculation, 0.1□mL of fungal preculture and 0.01□mL of bacterial preculture were transferred into each tube. The optimal fungal-bacterial combination was selected based on the resulting MCFAs concentration. To further assess the temporal dynamics of metabolite production, a kinetic experiment was performed using co-cultures of *N*. sp. FC1 with either *M. hexanoica* or *P. alactolyticus*. Time-course sampling was conducted destructively by harvesting three biological replicates per condition at each timepoint.

### Analytical methods

Headspace pressure and composition were analyzed to evaluate fungal proliferation. Headspace pressure was first measured using a pressure gauge (98766-89, Traceable). Gas composition was analyzed by gas chromatography equipped with a thermal conductivity detector (HP 5890, Hewlett Packard) and CTR I column (Alltech). Helium gas was used as the carrier gas. An isothermal method was applied with 50 °C oven temperature and 10 mins of run time.

All liquid samples were filtered through a 0.22□μm membrane filter and stored at - 20 °C before further analysis. Samples were thawed, centrifuged at 10,000 × g for 5 mins, and diluted using HPLC grade water prior to the analysis. Lactate and formate concentrations were quantified using high-performance liquid chromatography (ICS5000 Dionex, Thermo Scientific) with a UV-detector and Aminex HPX-87H column (Bio-Rad). The column was maintained at 50 °C. The mobile phase consisted of 5 mM sulfuric acid, and elution was performed at a flow rate of 0.6 mL/min. Data acquisition and analysis were carried out using Chromeleon 7 software (Thermo). For volatile fatty acids (acetate, propionate, butyrate, pentanoate, hexanoate, heptanoate, and octanoate) quantification, samples were acidified by adjusting formate concentration to 100 mM and were analyzed using gas chromatography (8900 GC System, Agilent) equipped with a mass spectrometry (7000D Triple Quadrupole MS, Agilent) and DB-FatWax column (Agilent). Helium gas was used as the carrier gas with 1 mL/min flow rate, and oven temperature gradient was set as follows: 80 °C for 1 min, a 20 °C/min ramp until 210 °C, 20 °C/min until 250 °C and finally held for 5 mins. The mass spectrometer was operated in dynamic multiple reaction monitoring (dMRM) mode. Target analytes were ionized and fragmented by electron ionization, and the precursor and product ions of target analytes were selected using Agilent software (MassHunter Optimizer, Agilent).

### Carbon mass balance experiments and conversion efficiency calculation

Carbon mass balance experiments were conducted under three conditions: (1) uninoculated control, (2) *N*. sp. FC1 monoculture, and (3) co-culture of *N*. sp. FC1 and *M. hexanoica*. All conditions were prepared in five biological replicates using Hungate tubes with a 10□mL working volume of M2 medium supplemented with reed canary grass (10 g/L) and yeast extract (1 g/L). Inoculated cultures were incubated for 7 days prior to mass balance analysis. Samples were analyzed for solid, aqueous, and gas fractions. Headspace pressure and gas composition was analyzed to quantify gaseous carbon dioxide. The entire culture content was then filtered using a vacuum filtration apparatus with a pre-weighed 0.22□μm membrane filter to separate the solid and liquid phases. The liquid fraction was collected in a 50□Ml Falcon tube, while the solid retained on the filter was dried at 105°C and weighed to determine the dry solid mass. Carbon content of the solid fraction was further analyzed via elemental analysis (Flash 2000, Thermo Scientific). The liquid fraction was analyzed for organic acid (i.e., lactate, formate and volatile fatty acids) concentrations. In addition, total organic carbon (TOC) and inorganic carbon (IC) were quantified using a TOC analyzer (TOC-L, Shimadzu). Lignocellulose-to-product conversion yield of *N*. sp. FC1 was calculated by dividing the total grams of carbon in fungal fermentation products (lactate, acetate, and formate) by the grams of carbon in the supplemented reed canary grass. The lignocellulose-to-MCFA conversion efficiency of co-cultures was calculated by dividing the total grams of MCFA carbon (hexanoate and octanoate) by the grams of carbon in the supplemented reed canary grass.

### Proteomics sample preparation and analysis

Proteomic samples were collected from both monoculture and co-culture conditions. There are three sources of protein in this experiment, which are *N*. sp. FC1, *M. hexanoica*, and grass used as the substrate. Sorghum was selected as an alternative lignocellulosic substrate for proteomic experiments due to the availability of fully annotated genome, which is necessary for the identification of potential plant-derived proteins. To validate the substitution of reed canary grass with sorghum, we compared fermentation kinetics and metabolite profiles of cultures grown on sorghum and reed canary grass under both monoculture and co-culture conditions. Fungal monocultures and fungal-bacterial co-cultures were inoculated in Hungate tubes with a 10□mL working volume of M2 medium supplemented with sorghum (10 g/L) and yeast extract (1 g/L). Bacterial monocultures were inoculated in the same condition but also supplemented with lactate, and acetate (1.8, and 2.6□g/L, respectively). No significant differences were observed in fermentation trends and yields between experiments using reed canary grass and those using sorghum (Supplementary Figure 4). Particularly, *N*. sp. FC1 rapidly produced its fermentation products within the first 5 days (Supplementary Figure 4A), and the co-culture of *N*. sp. FC1 and *M. hexanoica* showed higher hexanoate and octanoate production compared to the *M. hexanoica* monoculture (Supplementary Figure 4D). To minimize disruption of plant cell walls and release of sorghum proteins, cell lysis was performed using bead beating instead of cryo-grinding, resulting in sorghum-derived peptides comprising less than 0.1% of total spectra.

Biomass was harvested during the mid-exponential growth phase. Four fungal monocultures and three fungal-bacterial co-cultures were sampled after 60□h of incubation, while 12 bacterial monocultures were collected after 36□h. Solid samples were obtained by centrifugation, washed once with phosphate buffer saline (pH 7.0), and flash-frozen with liquid nitrogen. Cell lysis, protein isolation, and digestion were performed using procedures outlined by Chirania et al. [6], with minor modifications. Briefly, biomass was lysed by bead beating using 0.15□mm zirconium oxide beads in Tris-HCl buffer (100□mM, pH□8.0) containing 4% sodium dodecyl sulfate and 10□mM DL-Dithiothreitol. Lysates were cleared by centrifugation at 21,000□×□g for 10□min and heated at 90□°C for 10□min to denature proteins. Cysteine residues were blocked by alkylation with 30□mM iodoacetamide in the dark for 20□min at room temperature. Proteins were purified by chloroform-methanol extraction, washed with methanol, air-dried, and resolubilized in ammonium bicarbonate (ABC) buffer (100□mM, pH□8.0) containing 4% sodium deoxycholate (SDC). For bacterial monoculture samples, proteins from four tubes were pooled during resolubilization to ensure sufficient protein yield for downstream analysis, generating triplicate samples. Protein concentrations were determined using the BCA assay, and 250□μg of each protein sample was concentrated using 10□kDa MWCO centrifugal filters, rinsed with ABC buffer, and digested with MS-grade trypsin (1:75 w/w) overnight at room temperature, with an additional 3□h digestion after fresh trypsin addition. Tryptic peptides were recovered by filtering through 10□kDa MWCO centrifugal filters. The remaining SDC was removed by formic acid precipitation and water-saturated ethyl acetate extraction. Lastly, peptide samples were dried in a SpeedVac and resuspended in 0.5% formic acid.

Peptide separation, detection, and raw mass spectra data processing were performed as previously described by Gois et al. [35], with minor modifications. Peptide samples were loaded onto an EASY-nano LC 1000 system coupled to a Q-Exactive Orbitrap mass spectrometer (Thermo Scientific) and separated on a custom-packed SilicaTip emitter column (New Objective) using ReproSil-Pur C18-AQ resin (Dr. Maisch GmbH). Two solvent solutions comprised 0.1% formic acid in water (Solvent A) and 0.1% formic acid in acetonitrile (Solvent B) delivered at a constant flow rate of 250□nL/min. Chromatographic separation was achieved using the following linear gradient: 0 to 5 min 0% B, increase to 10% B; 5 to 93 min, increase to 40% B; 93 to 95 min, ramp to 95% B; held at 95% B until 105 min; returned to 0% B by 106 min and held until 120 min for column re-equilibration. The mass spectrometer was operated in positive ionization mode with a spray voltage of 3□kV and maintaining the capillary at 275□°C. Full MS scans were collected across a mass-to-charge range of 400-2,000□m/z at a resolution of 70,000 (automatic gain control target: 1e6; max injection time: 30□ms). For tandem mass spectrometry (MS/MS) acquisition, a Top10 data-dependent method was employed and selected the ten most abundant precursor ions within a 0.4□m/z isolation window. MS2 scans were acquired at a resolution of 17,500 (normalized collision energy: 27; max injection time: 50□ms).

Tandem mass spectra were processed with Global Proteome Machine pipeline (GPM; thegpm.org) pipeline (version 2020.11.12.1) and searched with X! Tandem (version X! Tandem Aspartate 2020.11.12.1) against custom protein databases for Sorghum, *N*. sp. FC1 and *M. hexanoica*. To construct a database, the genome assembly of *M. hexanoica* (GCF_012843505.1; ASM1284350v1) was annotated using Prokka (v1.14.6) and eggNOG-mapper (emapper v2.1.12, eggNOG database v5.0.2). The resulting genome annotation files are provided as Supplementary dataset 2. Database searches were performed with fixed carbamidomethylation (cysteine, selenocysteine) and variable modifications allowing for N-terminal pyro-glutamate formation (from glutamine, glutamic acid), N-terminal ammonia loss, deamidation (asparagine, glutamine), and oxidation or dioxidation events on methionine and tryptophan residues. Peptide and protein identifications were validated with Scaffold (v5.1.2, Proteome Software Inc.), using thresholds of greater than 95% probability for peptides and greater than 99% probability for proteins, with a requirement for at least two unique peptides. Protein probabilities were assigned by the Protein Prophet algorithm, while proteins containing shared peptides that could not be uniquely distinguished by MS/MS were grouped according to the principle of parsimony. Quantification was performed using the Top3 total ion current (TIC) approach, and spectral counts for fungal and bacterial proteins were normalized separately by scaling each organism’s total spectra within a sample to the average total spectra across all samples. Statistical significance was assessed using t-tests corrected with the Benjamini-Hochberg method. Additionally, identified CAZyme proteins were annotated using dbCAN3 (HMMER-based annotation against CAZy family profiles) and InterProScan (v5.75-106.0) to predict catalytic domains and carbohydrate-binding modules (CBMs).

### Technoeconomic analysis

Techno-economic analysis (TEA) was performed using BioSTEAM [39], an open-source Python platform for process simulation and cost estimation. This analysis aimed to assess whether fungal-bacterial co-cultures for direct conversion of reed canary grass into MCFAs could achieve economic viability and to identify key process parameters required for scale-up. The process configuration included four main stages: media preparation, fermentation, product separation, and distillation. BioSTEAM was used to simulate mass and energy flows for each unit operation, coupled with equipment design and cost algorithms to estimate capital and operating costs. Detailed assumptions, parameters, and sensitivity analyses are provided in Supplementary Note□2 and Supplementary Table□3.

### Outstanding Questions

▪ How can strain-level metabolic engineering be integrated with consortium-level cross-feeding to simultaneously maximize productivity and product selectivity in synthetic microbial systems?
▪ Which functional guilds should be incorporated into synthetic consortia to expand MCFA yield and robustness without increasing system complexity?
▪ How should bioreactors be designed to sustain in-line product extraction in high-solids, lignocellulose-rich systems?
▪ How can Design-Build-Test-Learn (DBTL) frameworks be coupled with automated, data-driven strain and consortium engineering to iteratively to accelerate optimization of integrated biomanufacturing process?

### Technology readiness

This study establishes an anaerobic fungal-bacterial consortium as a pretreatment-free platform for lignocellulose to medium-chain fatty acids conversion, along with its technoeconomic potential at laboratory scale. The Technology Readiness Level of our platform is approximately 4 based on demonstrated performance in batch fermentations with reed canary grass. As identified here via technoeconomic analysis, technology advancement requires reducing cultivation media costs and improving process intensification by operating at higher solids concentrations. Additionally, achieving higher MCFA yield would further improve technoeconomics, which can be achieved through metabolic engineering of the synthetic consortium and incorporating in-line MCFA extraction to reduce product toxicity.

## Author contributions

**BRK**: Conceptualization, Data curation, Formal analysis, Investigation, Methodology, Project administration, Software, Validation, Visualization, Writing – original draft, Writing – review & editing; **EMB**: Methodology and Resources for fungi cultivation, Project administration; **JPH**: Methodology and Resources for bacteria cultivation, Writing – review & editing; **IMG**: Methodology and Validation for differential proteomics, Writing – review & editing; **RF**: Data curation, Methodology, Software, Supervision, and Validation for differential proteomics; **TSL**: Investigation and Resources for isolating *Neocallimastix* sp. FC1; **JT, VB, SR**: Investigation, Methodology, and Resources for genomic DNA, total RNA extraction; **SM, JP, AL, JG, HH, RL, KB, IVG**: Investigation, Data curation, Methodology, Software, and Resources for genomic DNA, total RNA sequencing, genome annotation, and transcriptome assembly; **MAO’M**: Conceptualization, Funding acquisition, Methodology, Resources, Supervision; **CEL**: Conceptualization, Funding acquisition, Methodology, Resources, Supervision, Validation, Writing – review & editing.

## Data availability

The *Neocallimastix* sp. FC1 genome assembly and annotation are available from JGI MycoCosm portal [54] (https://mycocosm.jgi.doe.gov/NeoFC1_1) and have been deposited at DDBJ/EMBL/GenBank under the accession SRP609168. The mass spectrometry proteomics data have been deposited to the ProteomeXchange Consortium via the PRIDE partner repository with the dataset identifier PXD067542.

## Code availability

All codes used for techno-economic analysis of the lignocellulose-to-MCFAs conversion by fungal-bacterial consortia are available on Github (https://github.com/BK419/Technoeconomic_analysis_lignocellulose_to_MCFAs_fungal_bac terial_co-culture).

## Acknowledgements

We acknowledge funding support from the Department of Energy (DOE), Office of Science (DE-SC0022142), the Institute for Collaborative Biotechnologies and National Science Foundation (2128271). This work was also funded by the DOE Joint BioEnergy Institute (http://www.jbei.org) supported by the Office of Biological and Environmental Research of the DOE Office of Science through contract DE-AC02–05CH11231 between Lawrence Berkeley National Laboratory. We further acknowledge support from the Natural Sciences and Engineering Research Council of Canada (RGPIN-2021-02684, NSERC-CREATE 528163-201). The work (proposal: 10.46936/10.25585/60000510) conducted by the U.S. Department of Energy Joint Genome Institute (https://ror.org/04xm1d337), a DOE Office of Science User Facility, is supported by the Office of Science of the U.S. Department of Energy operated under Contract No. DE-AC02-05CH11231. We sincerely thank Dr. Cameron Strachan (University of Veterinary Medicine Vienna) for generously performing the genome annotation of *Megasphaera hexanoica* and providing guidance on annotation processing

