## SUPPLEMENTARY INFORMATION for "Engineering anaerobic fungal-bacterial consortia for direct conversion of lignocellulosic biomass into medium-chain fatty acids"

**Supplementary Note 1.** Nitrogen access limitation and adaptive metabolic shifts in *Megasphaera hexanoica* under co-culture

Under co-culture conditions, *M. hexanoica* exhibited distinct changes in protein abundance related to nitrogen metabolism (Supplementary Table 2). Notably, NAD-specific glutamate dehydrogenase, ornithine carbamoyltransferase, argininosuccinate synthase, and 2-aminoadipate transaminase were upregulated, indicating activation of the metabolic route that channels ammonia through glutamate, ornithine, and ultimately to arginine. This pathway plays a central role in assimilating inorganic nitrogen such as ammonia into organic intermediates. Glutamate serves as a primary nitrogen donor in biosynthesis, and its downstream conversion to ornithine and arginine supports intracellular nitrogen storage, buffering, and redistribution. The coordinated upregulation of these enzymes suggests that *M. hexanoica* actively reallocates internal nitrogen to sustain metabolic activity when access to extracellular nitrogen sources is constrained. Additionally, the downregulation of O-acetylserine sulfhydrylase points to a decrease in cysteine utilization. This metabolic adjustment likely reflects competition with the fungal partner for shared nitrogen pools such as ammonia, cysteine, and amino acids in yeast extract, and indicates a reorganization of nitrogen metabolism that enables *M. hexanoica* to maintain homeostasis under nutrient-constrained conditions.

**Supplementary Note 2.** Assumptions and parameters used in techno-economic analysis (TEA)

The modeled bioprocess consists of four main stages: media preparation, fermentation, separation, and distillation (Figure 5A). A fixed biomass input of 2000 t/day of grass was assumed, with an initial grass loading of 0.01 kg grass per kg medium. In the media preparation step, 30 % of the water was first mixed with grass, heated to 121°C, then cooled to 30°C before being combined with the remaining water with medium components. In the fermentation stage, the product yield was assumed to be 1 g MCFAs per 10 g grass, and either sole hexanoic acid or octanoic acid production was considered. The bioreactor system operated in 8 parallel batch reactors, each with a 7-day fermentation time and 1-day cleaning cycle. Following fermentation, solids and liquids were separated. The solid fraction was directed to a boiler-turbogenerator system for combustion and energy recovery, while the liquid fraction underwent liquid-liquid extraction using 2-octanol as the solvent. The solvent-to-product ratio was set at 4:1 (mass basis), and extraction efficiencies were assumed to be 96 % for hexanoic acid and 99 % for octanoic acid^1^. A 5 % solvent loss to the wastewater stream was assumed, with a wastewater treatment cost of 0.31 USD/m³^2^. The MCFAs-enriched extractant was further processed via distillation, yielding a final product purity of 99 %. The 2-octanol solvent was recycled for subsequent extraction cycles. Material costs and unit prices are provided in Supplementary Table 3. This design adopts a conservative approach, as more efficient separation methods, such as membrane-based extraction, could be considered in future analyses. Additionally, the residual lignocellulosic solids could potentially be sold as dried solid fuel to enhance economic viability.

The initial TEA indicated that the high medium cost (127.2 USD/L) was the primary economic bottleneck. A preliminary scenario assuming 0.1 kg_MCFAs/kg_grass_yield and 0.01 kg_grass/kg_medium_loading resulted in an estimated profit of only 3–5 USD/L_medium_loading, which is economically unfeasible. However, the assumed grass loading of 0.01 kg/kg is not realistic for commercial-scale systems, and the use of costly components such as cysteine, buffering agents, and yeast extract is impractical. To identify key process parameters that influence economic performance, we performed a sensitivity analysis using Spearman correlation between input variables and model outputs (IRR and utility cost). Seven parameters were evaluated: medium cost, MCFAs yield, RCG (reed canary grass) input, fermentation time, octanol price, RCG price, and wastewater treatment cost. As shown in Fig. 5c-d, medium cost, MCFAs yield, and RCG input exhibited the strongest correlations with IRR, while RCG input and MCFAs yield were the most influential factors for utility cost. These findings highlight the importance of minimizing medium cost and maximizing biomass loading and product yield to improve process profitability. All tested parameter ranges and baseline values are provided in Supplementary Table 3. To identify the threshold medium cost (USD/kg MCFAs) required for profitability, we defined a new metric by directly subtracting medium cost from the MCFAs market price. In this analysis, we used a more realistic grass loading of 0.1 kg/kg. We then calculated the combinations of medium cost and MCFAs yield that would result in target internal rates of return (IRR = 0.1, 0.15, 0.2). The range of medium cost was varied from 0 to the MCFAs price, and MCFAs yield was explored from 0.01 to its theoretical maximum. The theoretical maximum yield was calculated assuming 35.2 % and 28.0 % of cellulose and hemicellulose content in grass and 100 % conversion efficiency into hexanoate or octanoate.

**Supplementary Figures**

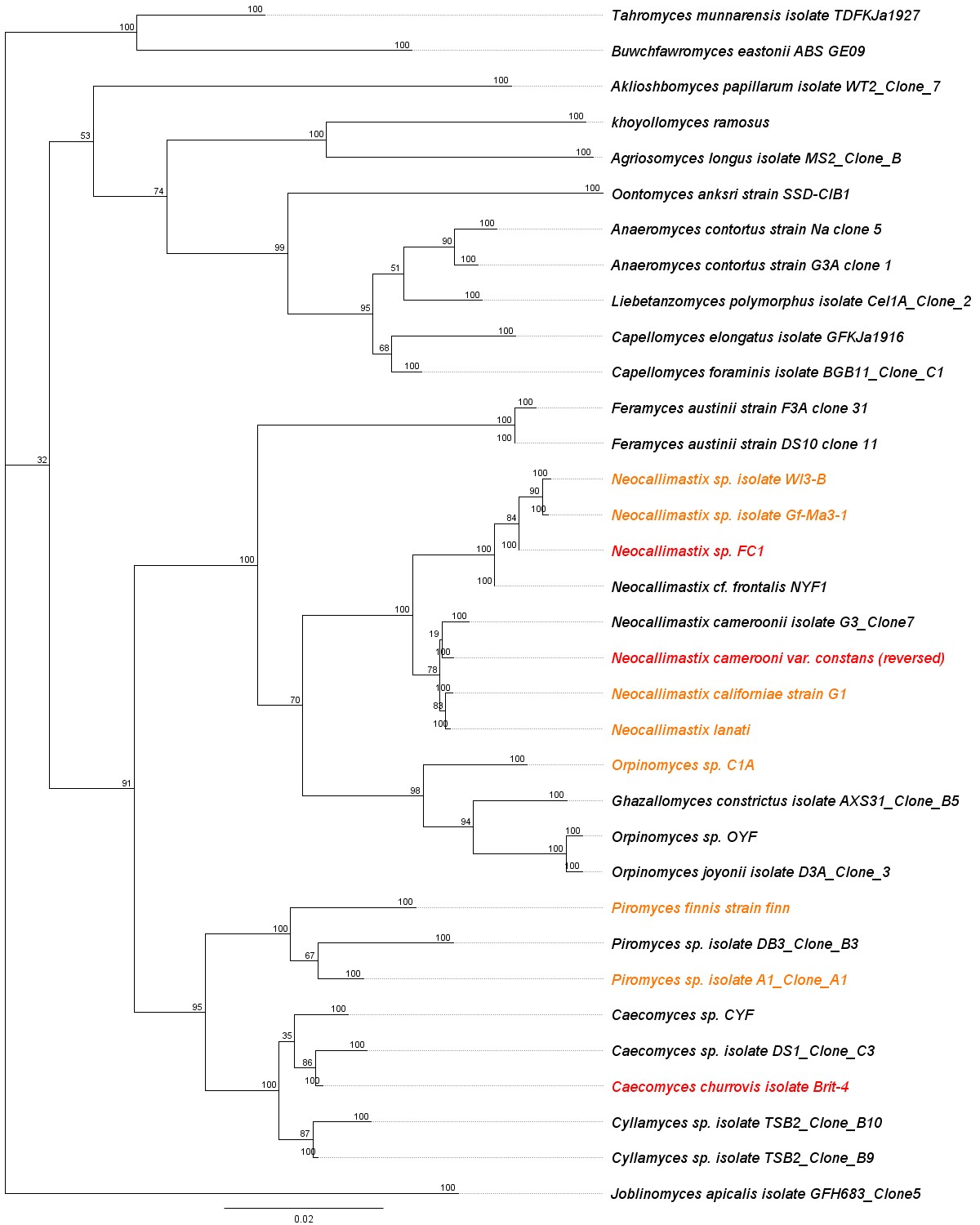

**Supplementary Figure 1.** Phylogenetic tree of anaerobic fungi based on LSU rRNA gene sequences. Maximum likelihood phylogenetic tree of anaerobic fungi constructed using LSU rRNA gene sequences. Strains with fully annotated genomes are highlighted in yellow and red. Red labels indicate strains used in this study, while yellow labels represent other anaerobic fungi with publicly available genome annotations. Bootstrap values (shown at the nodes) indicate the level of support for each clade.

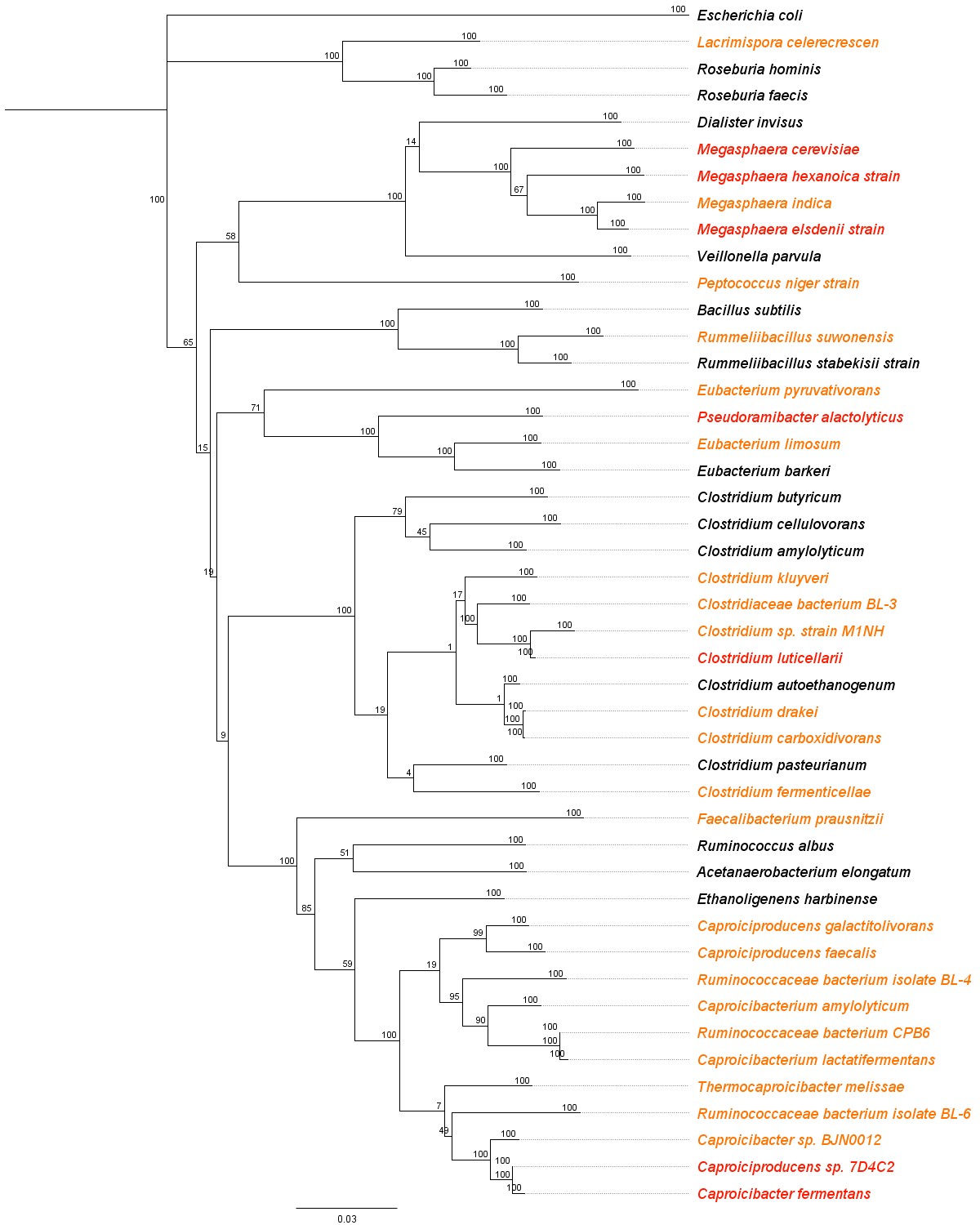

**Supplementary Figure 2.** Phylogenetic tree of bacterial chain elongators based on full-length 16S rRNA gene sequences. Maximum likelihood phylogenetic tree of bacterial strains constructed using full-length 16S rRNA gene sequences. Strains known to perform chain elongation are highlighted in yellow and red. Red-labeled strains were used in this study, while yellow-labeled strains represent other reported chain elongators. Bootstrap values (indicated at nodes) represent the statistical support for each branch.

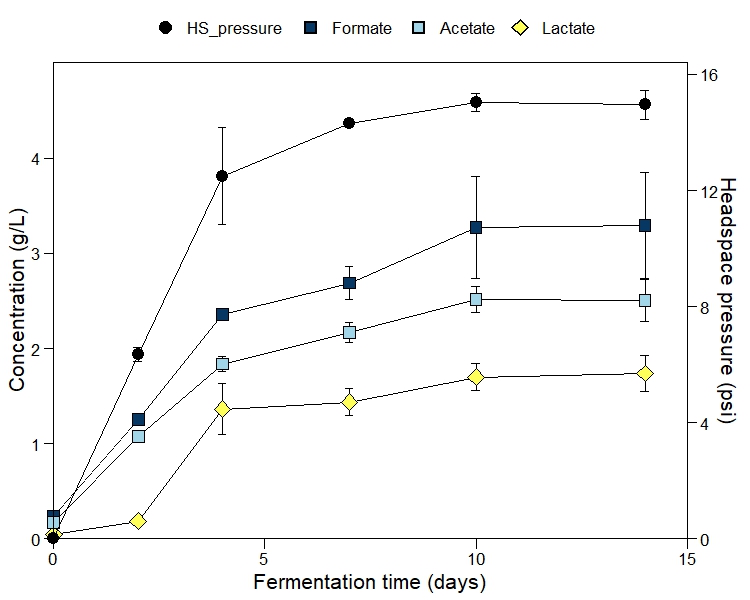

**Supplementary Figure 3.** Time-course profiles of gas and metabolite production in monoculture fermentation of *Neocallimastix* sp. FC1. Fungal monocultures were monitored for 14 days, and concentrations of formate, acetate, and lactate were quantified alongside headspace pressure measurements. All data represent the average of biological triplicates, with error bars indicating standard deviations.

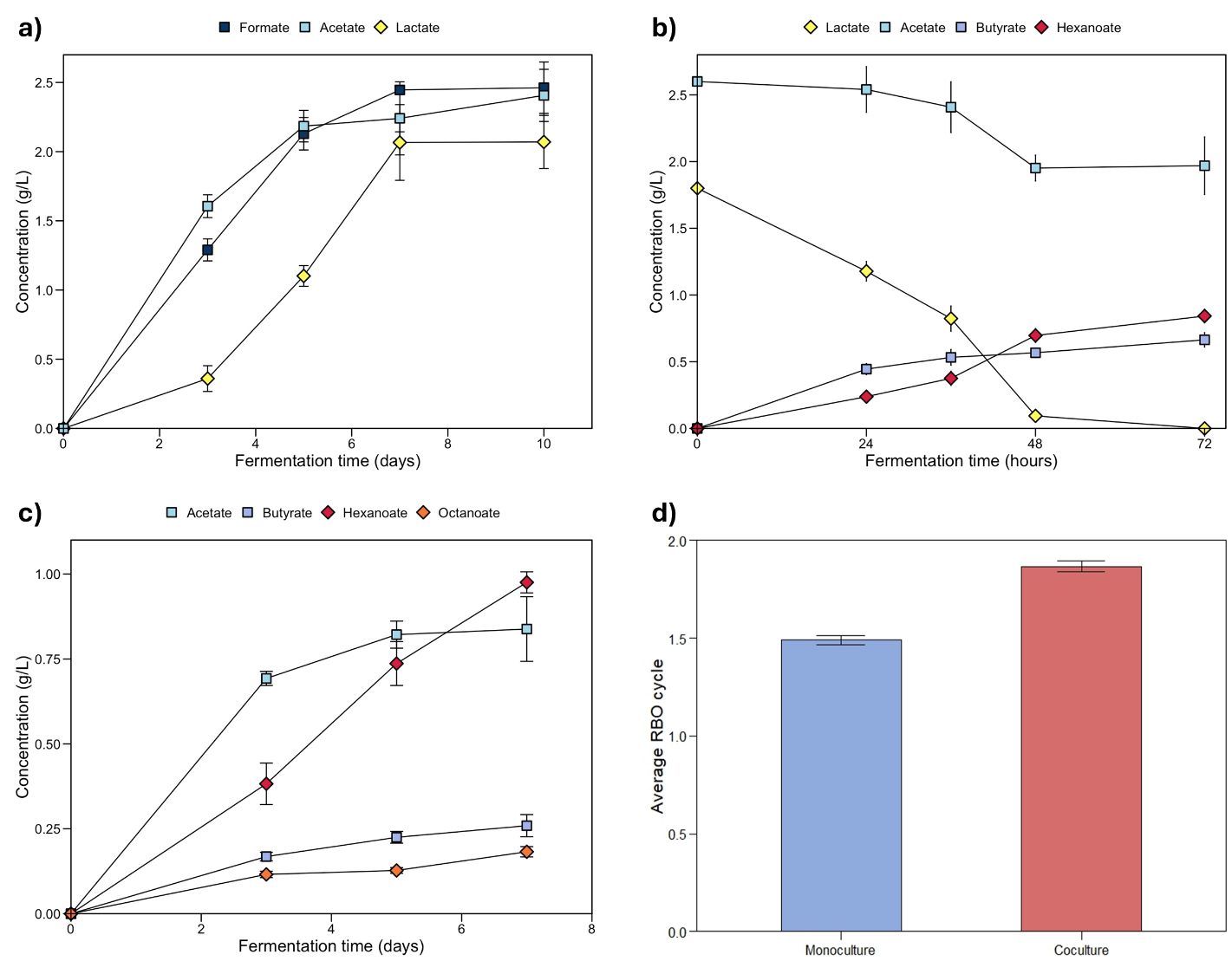

**Supplementary Figure 4.** Fermentation kinetics and average reverse β-oxidation (RBO) cycle in *Neocallimastix* sp. FC1 and *Megasphaera hexanoica* monoculture and co-culture systems using sorghum as the lignocellulosic substrate. **a-c**, Fermentation kinetics of *N.* sp. FC1 monoculture (**a**), *M. hexanoica* monoculture (**b**), and co-culture (**c**). **d**, Average RBO cycle in *M. hexanoica* monoculture and *N.* sp. FC1 and *M. hexanoica* co-culture. All data represent the average of biological triplicates, with error bars indicating standard deviations.

**Supplementary Tables**

**Supplementary Table 1.** Summary of the features of the genome of *Neocallimastix*. sp. FC1

| **Features** | **Value** |
| --- | --- |
| Genome size (Mbp) | 205.51 |
| No. of contigs | 1360 |
| No. of predicted genes | 30176 |
| No. of CAZymes (carbohydrate active enzyme) | 1453 |
| No. of GH (glycoside hydrolase) genes | 806 |
| No. of GH genes containing a fungal dockerin domain | 99 |

**Supplementary Table 2.** Spectral counts, significance scores, and fold changes of selected enzymes in *Megasphaera hexanoica* and *Neocallimastix* sp. FC1 proteome under monoculture and co-culture conditions. Monoculture and co-culture columns indicate Top 3 total ion current from each condition. Significance score refers to the negative base-10 logarithm of the p-value (-log_10_(p-value)) derived from differential protein abundance analysis. Fold change represents the absolute log_2_-transformed ratio of spectral counts between co-culture and monoculture (log_2_(Co-culture/Monoculture))). A dash (-) indicates cases where the protein was not detected in co-culture. Enzyme numbers listed before the enzyme names correspond to those shown in Fig. 3a; proteins without numbers are not included in the figure. Bold indicates enzymes with statistically significant differences in abundance between monoculture and co-culture (FDR < 0.05).

| ***Megasphaera hexanoica* proteins** | | | | | |
| --- | --- | --- | --- | --- | --- |
| **Enzyme** | **Accession number** | **Mono**  **culture** | **Co-culture** | **Significance score** | **Fold change** |
| 26. Lactate racemase | BOHMIINL_01620 | 8.1.E+07 | 8.9.E+07 | 0.84 | 0.1 |
| 27. L-lactate dehydrogenase | BOHMIINL_02481 | 5.3.E+06 | 1.4.E+07 | 0.32 | 1.4 |
|  | BOHMIINL_02311 | 9.6.E+06 | - | - | - |
| **28. Pyruvate:ferredoxin oxidoreductase** | **BOHMIINL_02337** | **3.9.E+08** | **6.1.E+08** | **2.57** | **0.6** |
|  | BOHMIINL_00087 | 3.2.E+07 | 5.2.E+07 | 0.61 | 0.7 |
|  | BOHMIINL_00223 | 2.9.E+07 | 3.8.E+07 | 0.13 | 0.4 |
|  | BOHMIINL_02317 | 5.8.E+06 | 3.3.E+07 | 1.16 | 2.5 |
|  | BOHMIINL_02456 | 2.6.E+06 | - | - | - |
| 29. Acetyl-CoA acetyltransferase | BOHMIINL_01501 | 3.5.E+08 | 2.9.E+08 | 0.66 | -0.3 |
| **30. 3-hydroxybutyryl-CoA dehydrogenase** | **BOHMIINL_00848** | **2.4.E+08** | **3.3.E+08** | **1.82** | **0.4** |
| **31. Enoyl-CoA hydratase** | **BOHMIINL_00849** | **2.0.E+08** | **1.2.E+09** | **4.35** | **2.6** |
| 32. Electron transfer flavoprotein | BOHMIINL_00532 | 9.6.E+08 | 1.1.E+09 | 0.46 | 0.2 |
|  | BOHMIINL_00533 | 1.5.E+08 | 1.8.E+08 | 0.48 | 0.3 |
|  | BOHMIINL_02216 | 3.1.E+06 | - | - | - |
|  | BOHMIINL_02217 | 1.3.E+06 | - | - | - |
| **33. Acyl-CoA dehydrogenase** | BOHMIINL_00555 | 2.7.E+08 | 5.0.E+08 | 1.12 | 0.9 |
|  | **BOHMIINL_00534** | **7.7.E+07** | **1.7.E+08** | **1.85** | **1.1** |
|  | BOHMIINL_02285 | 4.7.E+06 | - | - | - |
| 34. Butyryl-CoA:acetate CoA-transferase | BOHMIINL_00232 | 6.6.E+07 | 4.4.E+07 | 0.81 | -0.6 |
|  | BOHMIINL_02095 | 1.6.E+06 | - | - | - |
| 35. Phosphate acyltransferase | BOHMIINL_01032 | 2.9.E+07 | 4.3.E+07 | 0.76 | 0.6 |
|  | BOHMIINL_00318 | 7.4.E+05 | - | - | - |
| 36. Na(+)-translocating ferredoxin:NAD(+) oxidoreductase | BOHMIINL_01861 | 3.6.E+07 | 4.9.E+07 | 0.23 | 0.5 |
|  | BOHMIINL_01864 | 1.2.E+07 | - | - | - |
| 37. Proton-translocating ferredoxin:NAD(+) oxidoreductase | BOHMIINL_01859 | 1.5.E+07 | 3.2.E+07 | 0.91 | 1.1 |
|  | BOHMIINL_01860 | 5.5.E+06 | - | - | - |
| 38. ATP synthase | BOHMIINL_00806 | 1.2.E+07 | 3.6.E+07 | 1.64 | 1.6 |
|  | BOHMIINL_00804 | 5.2.E+06 | 6.7.E+06 | 0.08 | 0.4 |
|  | BOHMIINL_00805 | 3.7.E+06 | - | - | - |
|  | BOHMIINL_00802 | 1.2.E+07 | - | - | - |
|  | BOHMIINL_00803 | 9.9.E+05 | - | - | - |
| 39. Iron hydrogenase | BOHMIINL_02270 | 1.4.E+07 | 4.7.E+06 | 1.14 | -1.6 |
|  | BOHMIINL_01073 | 1.9.E+07 | 2.3.E+07 | 0.08 | 0.2 |
| Acetate kinase | BOHMIINL_00396 | 8.0.E+06 | - | - | - |
| 6,7-dimethyl-8-ribityllumazine synthase | BOHMIINL_01480 | 4.0.E+07 | 1.6.E+07 | 2.00 | -1.4 |
| Argininosuccinate synthase | BOHMIINL_00294 | 9.0E+07 | 1.7E+08 | 2.47 | 1.0 |
| NAD-specific glutamate dehydrogenase | BOHMIINL_01168 | 3.2E+05 | 4.4E+07 | 4.67 | 7.1 |
| 2-aminoadipate transaminase | BOHMIINL_00416 | 5.8E+06 | 9.4E+07 | 4.07 | 4.0 |
| Ornithine carbamoyltransferase | BOHMIINL_00422 | 2.2E+07 | 5.2E+07 | 2.01 | 1.2 |
| O-acetylserine sulfhydrylase | BOHMIINL_01018 | 1.4E+07 | 1.6E+06 | 2.58 | -3.1 |
| ***Neocallimastix* sp. FC1 proteins** | | | | | |
| **Enzyme** | **Accession number** | **Mono**  **culture** | **Co-culture** | **Significance score** | **Fold change** |
| 2. Hexokinase | e_gw1.132.99.1 | 4.6E+07 | 5.4E+07 | 0.50 | 0.2 |
|  | fgenesh1_pm.57_#_8 | 7.3E+07 | 6.1E+07 | 0.44 | -0.3 |
|  | fgenesh2_ kg.642___29___ TRINITY_DN1408_ c0_g3_i1 | 6.0E+07 | 5.4E+07 | 0.27 | -0.1 |
|  | e_gw1.252.31.1 | 7.2E+07 | 3.5E+07 | 0.59 | -1.0 |
|  | fgenesh2_ kg.122___83___ OBPOO_transcript/5798 | 3.4E+07 | 3.0E+07 | 0.24 | -0.2 |
| 3. Glucose-6-phosphate isomerase | fgenesh2_ kg.147___92___ TRINITY_DN608_ c0_g1_i5 | 3.1E+07 | 2.4E+07 | 0.16 | -0.4 |
| 4. 6-phosphofructokinase | e_gw1.13.301.1 | 3.0E+07 | 2.6E+07 | 0.14 | -0.2 |
|  | estExt_ Genewise1Plus.C_ 1600044 | 7.2E+06 | 9.2E+06 | 0.52 | 0.4 |
| 5. Fructose-bisphosphate aldolase | fgenesh2_ kg.136___204___ OBPOO_transcript/18767 | 1.7E+08 | 2.5E+08 | 0.86 | 0.6 |
|  | fgenesh2_ kg.152___553___ OBPOO_transcript/14027 | 1.7E+08 | 3.0E+08 | 1.32 | 0.8 |
| 6. Triosephosphate isomerase | fgenesh2_ kg.105___236___ TRINITY_DN2887_ c0_g1_i2 | 1.3E+08 | 1.3E+08 | 0.01 | 0.0 |
|  | fgenesh2_ kg.931___23___ TRINITY_DN2887_ c0_g1_i5 | 1.3E+08 | 1.3E+08 | 0.01 | 0.0 |
| 7. Xylose isomerase | fgenesh2_ kg.30___400___ OBPOO_transcript/8332 | 3.8E+08 | 2.6E+08 | 2.79 | -0.6 |
|  | fgenesh2_ kg.650___124___ OBPOO_transcript/13899 | 3.8E+08 | 2.5E+08 | 2.56 | -0.6 |
|  | fgenesh2_ kg.650___52___ OBPOO_transcript/7105 | 3.8E+08 | 2.5E+08 | 2.56 | -0.6 |
|  | fgenesh2_ kg.30___508___ OBPOO_transcript/6681 | 3.7E+08 | 2.5E+08 | 1.76 | -0.5 |
|  | fgenesh2_ kg.1196___30___ OBPOO_transcript/9060 | 3.1E+08 | 2.2E+08 | 1.40 | -0.5 |
|  | fgenesh2_ kg.197___50___ OBPOO_transcript/11442 | 2.7E+08 | 1.6E+08 | 0.82 | -0.7 |
| 8. D-xylulokinase | fgenesh2_ kg.30___449___ OBPOO_transcript/8542 | 5.9E+07 | 9.6E+07 | 1.15 | 0.7 |
|  | fgenesh2_ kg.768___64___ TRINITY_DN2724_ c0_g1_i4 | 7.1E+07 | 7.3E+07 | 0.22 | 0.0 |
| 9. Transketolase | gm4.16474_g | 5.7E+07 | 8.7E+07 | 0.58 | 0.6 |
|  | gm4.35776_g | 1.4E+07 | 2.6E+07 | 1.10 | 0.9 |
| 10. Glyceraldehyde-3-phosphate dehydrogenase 3 | fgenesh2_ kg.138___66___ OBPOO_transcript/18269 | 7.2E+07 | 7.4E+07 | 0.15 | 0.1 |
| 11. Phosphoglycerate kinase | fgenesh2_ kg.790___1___ TRINITY_DN3976_ c0_g1_i24 | 1.1E+08 | 1.2E+08 | 0.30 | 0.2 |
|  | fgenesh2_ kg.9___198___ OBPOO_transcript/11579 | 9.9E+07 | 9.6E+07 | 0.05 | -0.1 |
| 12. Phosphoglycerate mutase | fgenesh2_ kg.52___399___ TRINITY_DN553_ c0_g1_i1 | 2.5E+07 | - | - | - |
|  | fgenesh2_ kg.852___11___ OBPOO_transcript/7816 | 4.3E+07 | 3.1E+07 | 0.32 | -0.5 |
| 13. Enolase | fgenesh1_pg.554_#_14 | 4.5E+08 | 3.9E+08 | 0.38 | -0.2 |
| 14. Pyruvate kinase | estExt_ Genewise1Plus.C_ 460152 | 2.2E+07 | 1.4E+07 | 1.19 | -0.6 |
| 15. Phosphoenolpyruvate carboxykinase | fgenesh2_ kg.422___118___ OBPOO_transcript/2067 (+1) | 5.0E+08 | 5.7E+08 | 0.22 | 0.2 |
|  | fgenesh1_pm.460_#_3 | 3.7E+07 | 5.4E+07 | 0.39 | 0.5 |
|  | fgenesh2_ kg.148___151___ TRINITY_DN11490_ c0_g1_i1 (+3) | 4.1E+06 | - | - | - |
| 16. Malate dehydrogenase | fgenesh2_ kg.861___147___ OBPOO_transcript/18110 | 4.0E+08 | 4.2E+08 | 0.10 | 0.1 |
|  | e_gw1.77.82.1 | 2.7E+07 | 2.8E+07 | 0.10 | 0.1 |
|  | fgenesh2_ kg.290___115___ OBPOO_transcript/22039 | 1.6E+07 | 2.9E+07 | 0.46 | 0.8 |
|  | fgenesh2_ kg.54___360___ OBPOO_transcript/21314 | 3.4E+08 | 3.9E+08 | 0.35 | 0.2 |
| 17. Malic enzyme | fgenesh2_ kg.315___50___ TRINITY_DN1051_ c0_g1_i11 | 7.4E+08 | 6.2E+08 | 0.58 | -0.3 |
|  | fgenesh2_ kg.267___15___ OBPOO_transcript/2566 | 7.4E+08 | 6.2E+08 | 0.58 | -0.3 |
|  | fgenesh2_ kg.67___211___ TRINITY_DN1051_ c0_g1_i5 | 7.4E+08 | 6.2E+08 | 0.58 | -0.3 |
|  | fgenesh2_ kg.75___113___ OBPOO_transcript/1828 | 6.9E+08 | 5.8E+08 | 0.93 | -0.2 |
| 18. Pyruvate formate lyase | fgenesh2_ \|kg.25___538___ OBPOO_transcript/1170 (+1) | 3.4E+08 | 4.0E+08 | 0.54 | 0.2 |
|  | fgenesh2_ kg.74___290___ OBPOO_transcript/1018 | 2.1E+08 | 2.5E+08 | 0.52 | 0.2 |
|  | fgenesh2_ kg.165___250___ OBPOO_transcript/1013 | 2.1E+08 | 2.5E+08 | 0.58 | 0.3 |
|  | fgenesh2_ kg.167___162___ OBPOO_transcript/1018 (+1) | 2.0E+08 | 2.5E+08 | 0.59 | 0.3 |
|  | fgenesh2_ kg.645___76___ OBPOO_transcript/1701 | 1.9E+08 | 2.4E+08 | 0.70 | 0.3 |
|  | gm4.7561_g | 1.7E+08 | 1.9E+08 | 0.39 | 0.2 |
|  | fgenesh2_ kg.361___30___ OBPOO_transcript/2895 | 1.4E+08 | 1.7E+08 | 0.45 | 0.3 |
|  | fgenesh2_ kg.639___19___ OBPOO_transcript/1509 | 2.1E+08 | 1.6E+08 | 0.31 | -0.4 |
|  | fgenesh2_ kg.116___69___ OBPOO_transcript/631 | 3.7E+07 | 5.1E+07 | 0.56 | 0.5 |
|  | fgenesh2_ kg.116___152___ OBPOO_transcript/631 | 3.3E+07 | 4.3E+07 | 0.28 | 0.4 |
|  | estExt_fgenesh1_ pg.C_4090002 | 2.7E+07 | 3.8E+07 | 0.34 | 0.5 |
|  | estExt_Genemark4.C_ 2310030 | 2.7E+07 | 2.3E+07 | 0.45 | -0.2 |
|  | fgenesh2_ kg.582___58___ OBPOO_transcript/631 | 3.5E+07 | 2.3E+07 | 0.50 | -0.6 |
|  | fgenesh2_ kg.238___8___ OBPOO_transcript/22160 | 1.3E+07 | 1.6E+07 | 0.34 | 0.3 |
| 19. Acetyl-CoA hydrolase-succinyl-CoA-acetyl-CoA transferase | fgenesh2_ kg.368___19___ OBPOO_transcript/5328 | 8.4E+07 | 1.1E+08 | 0.52 | 0.4 |
| 20. Succinyl-CoA synthetase | e_gw1.229.6.1 | 3.5E+08 | 2.5E+08 | 0.68 | -0.5 |
|  | fgenesh2_ kg.248___91___ OBPOO_transcript/21822 (+1) | 2.2E+08 | 2.4E+08 | 0.11 | 0.1 |
|  | fgenesh2_ kg.85___100___ OBPOO_transcript/19694 | 1.2E+08 | 1.7E+08 | 1.17 | 0.5 |
| 21. Citrate synthase | e_gw1.637.6.1 | 6.7E+06 | 6.0E+06 | 0.11 | -0.2 |
| 22. Aconitate hydratase | fgenesh2_ kg.206___16___ TRINITY_DN2085_ c0_g2_i1 | 3.1E+07 | 2.8E+07 | 0.09 | -0.1 |
|  | e_gw1.1156.6.1 | 9.8E+06 | 1.1E+07 | 0.29 | 0.2 |
| 23. Isocitrate dehydrogenase | fgenesh2_ kg.403___38___ TRINITY_DN3391_ c0_g1_i4 | 1.1E+08 | 1.9E+08 | 1.49 | 0.8 |
|  | fgenesh2_ kg.40___73___ TRINITY_DN3346_ c0_g1_i1 | 3.4E+07 | 2.0E+07 | 2.81 | -0.7 |
| 24. D-lactate dehydrogenase | fgenesh2_ kg.995___4___ TRINITY_DN1005_ c1_g1_i1 | 1.1E+07 | 1.5E+07 | 0.48 | 0.4 |
|  | fgenesh2_ kg.654___5___ TRINITY_DN1005_ c1_g1_i1 | 9.0E+06 | 1.4E+07 | 0.88 | 0.7 |
| 25. Alcohol dehydrogenase | fgenesh2_ kg.80___286___ OBPOO_transcript/14856 | 1.2E+08 | 1.7E+08 | 1.16 | 0.5 |
| 25. Aldehyde reductase | e_gw1.564.23.1 | 1.4E+06 | 4.6E+05 | 0.37 | -1.6 |
| 25. NADP-dependent alcohol dehydrogenase | fgenesh2_ kg.153___15___ TRINITY_DN4270_ c0_g1_i2 | 1.8E+07 | 1.5E+07 | 0.43 | -0.3 |

**Supplementary Table 3.** Assumed material costs, product prices, and parameter ranges or values used in the techno-economic analysis (TEA)

| **Material** | **Price (USD/kg)** | **Reference** | | | |
| --- | --- | --- | --- | --- | --- |
| Water | 0.000353 | Choi et al., 2024^2^ | | | |
| Reed canary grass | 0.075 | Lynd et al., 2022^3^ | | | |
| NaHCO_3_ | 0.8 | https://catcost.chemcatbio.org/materials-library | | | |
| Yeast extract | 50.1 | Choi et al., 2024^2^ | | | |
| KH_2_PO_4_ | 2.77 | Choi et al., 2024^2^ | | | |
| (NH_4_)_2_SO_4_ | 0.25 | https://catcost.chemcatbio.org/materials-library | | | |
| NaCl | 0.46 | Choi et al., 2024^2^ | | | |
| MgSO_4_ | 0.67 | https://catcost.chemcatbio.org/materials-library | | | |
| CaCl_2_ | 0.47 | https://catcost.chemcatbio.org/materials-library | | | |
| K_2_HPO_4_^1)^ | 2.75 | Debergh et al., 2022^4^ | | | |
| L-cysteine^2)^ | 63.2 | https://www.chemicalbook.com/ProductList_En.aspx?kwd=cysteine&a=United%20States&left=True&c=5KG%7C10KG%7C25KG#J_Condition | | | |
| Thiamine | 520 | Choi et al., 2024^2^ | | | |
| Riboflavin^1)^ | 110 | Debergh et al., 2022^4^ | | | |
| Calcium  D-pantothenate^1)^ | 110 | Debergh et al., 2022^4^ | | | |
| Nicotinic acid^1)^ | 110 | Debergh et al., 2022^4^ | | | |
| Folic acid^1)^ | 110 | Debergh et al., 2022^4^ | | | |
| Cyanobalamin^1)^ | 110 | Debergh et al., 2022^4^ | | | |
| Biotin^1)^ | 110 | Debergh et al., 2022^4^ | | | |
| Pyridoxin^1)^ | 110 | Debergh et al., 2022^4^ | | | |
| $\rho$-Aminobenzoic acid^1)^ | 110 | Debergh et al., 2022^4^ | | | |
| MnCl_2_^1)^ | 27.5 | Debergh et al., 2022^4^ | | | |
| NiCl_2_^1)^ | 27.5 | Debergh et al., 2022^4^ | | | |
| Na_2_MoO_4_$\cdot$2H_2_O^1)^ | 27.5 | Debergh et al., 2022^4^ | | | |
| H_3_BO_3_ | 1.41 | https://catcost.chemcatbio.org/materials-library | | | |
| FeSO_4_$\cdot$H_2_O | 1.87 | https://catcost.chemcatbio.org/materials-library | | | |
| CoCl_2_^1)^ | 27.5 | Debergh et al., 2022^4^ | | | |
| SeO_2_^1)^ | 27.5 | Debergh et al., 2022^4^ | | | |
| NaVO_3_^1)^ | 27.5 | Debergh et al., 2022^4^ | | | |
| ZnCl_2_ | 1.31 | https://catcost.chemcatbio.org/materials-library | | | |
| CuCl_2_^1)^ | 27.5 | Debergh et al., 2022^4^ | | | |
| HCl | 0.13 | https://catcost.chemcatbio.org/materials-library | | | |
| Hemin^2)^ | 1700 | https://www.chemicalbook.com/ProductList_En.aspx?cbn=CB2409023&c=500g&left=True#J_Condition | | | |
| NaOH | 0.97 | https://catcost.chemcatbio.org/materials-library | | | |
| Ethanol | 1.16 | https://catcost.chemcatbio.org/materials-library | | | |
| Hexanoic acid^3)^ | 3.795 | Dessi et al., 2021^5^ | | | |
| Octanoic acid | 5 | Scarborough et al. 2018^1^ | | | |
| 2-Octanol | 1.55 | Scarborough et al. 2018^1^ | | | |
| **Parameter** | **Unit** | **Lower boundary** | **Most probable value** | **Upper boundary** | **Spearman's**  **correlation**  **with IRR** |
| Medium cost | USD/ kg_MCFA | 0 | - | 3.5 | -0.434 |
| MCFAs yield | kg_MCFA/kg_RCG | 0.1 | - | 0.41 | 0.224 |
| RCG (reed canary grass) input | kg_RCG/ kg_medium | 0.01 | - | 0.1 | 0.293 |
| Fermentation time | Day | 1 | 7 | 10 | -0.010 |
| Octanol price | USD/kg | 0.5 | 1.5 | 2.5 | -0.135 |
| RCG price | USD/kg | 0.03 | 0.06 | 0.12 | -0.033 |
| Wastewater treatment cost | USD/L | 0 | 0.00031 | 0.0031 | -0.029 |
| **Parameter** | **Unit** | **Value** | **Reference** | | |
| Chemical engineering plant  cost index (CEPCI) | - | 800.2 | 2024, May  https://toweringskills.com/financial-  analysis/cost-indices/ | | |
| Electricity price | USD/ kWh | 0.1 | https://www.eia.gov/electricity/monthly/epm_table_grapher.php?t=epmt_5_6_a | | |
| Heat transfer efficiency | - | 0.9 | https://biosteam.readthedocs.io/en/latest/tutorial/Building_a_biorefinery.html | | |
| Boiler efficiency | - | 0.8 | https://biosteam.readthedocs.io/en/latest/tutorial/Building_a_biorefinery.html | | |
| Turbogenerator efficiency | - | 0.85 | https://biosteam.readthedocs.io/en/latest/tutorial/Building_a_biorefinery.html | | |
| Steam utility temperature | K | 529.2 | https://biosteam.readthedocs.io/en/latest/tutorial/Building_a_biorefinery.html | | |
| Steam utility pressure | MPa | 0.44 | https://biosteam.readthedocs.io/en/latest/tutorial/Building_a_biorefinery.html | | |
| Makeup water price | USD/kg | 0.000254 | https://biosteam.readthedocs.io/en/latest/tutorial/Building_a_biorefinery.html | | |
| Cooling water regeneration | USD/kg | 0 | https://biosteam.readthedocs.io/en/latest/tutorial/Building_a_biorefinery.html | | |
| Chilled water heat transfer price | USD/kg | 0 | https://biosteam.readthedocs.io/en/latest/tutorial/Building_a_biorefinery.html | | |

^1)^ The prices reported by Debergh et al. (2022) were in euros, and we applied an exchange rate of 1.1 USD/EUR.

^2)^ Assuming the facility operates in Canada, we set to the lowest prices available from suppliers in Canada and the United States.

^3)^ Dessi et al. (2021) reported a hexanoic acid price range of 2.7–4.2 EUR/kg. We used 3.45 EUR/kg, calculated as the midpoint of this range, and applied an exchange rate of 1.1 USD/EUR.
